# Single-cell sequencing of developing human gut reveals transcriptional links to childhood Crohn’s disease

**DOI:** 10.1101/2020.02.06.937110

**Authors:** Rasa Elmentaite, Alexander Ross, Kylie R. James, Daniel Ortmann, Tomas Gomes, Kenny Roberts, Komal Nayak, Liz Tuck, Omer Ali Bayraktar, Robert Heuschkel, Ludovic Vallier, Sarah A. Teichmann, Matthias Zilbauer

## Abstract

Human gut development requires the orchestrated interaction of various differentiating cell types. Here we generate an in-depth single-cell map of the developing human intestine at 6–10 weeks post-conception, a period marked by crypt-villus formation. Our analysis reveals the transcriptional profile of cycling epithelial precursor cells, which are distinct from LGR5-expressing cells. We use computational analyses to show that these cells contribute to differentiated cell subsets directly and indirectly via the generation of LGR5-expressing stem cells and receive signals from the surrounding mesenchymal cells. Furthermore, we draw parallels between the transcriptomes of *ex vivo* tissues and *in vitro* fetal organoids, revealing the maturation of organoid cultures in a dish. Lastly, we compare scRNAseq profiles from paediatric Crohn’s disease epithelium alongside matched healthy controls to reveal disease associated changes in epithelial composition. Contrasting these with the fetal profiles reveals re-activation of fetal transcription factors in Crohn’s disease epithelium. Our study provides a unique resource, available at **www.gutcellatlas.org**, and underscores the importance of unravelling fetal development in understanding disease.

## Introduction

Development of the human intestine is a highly complex process that requires synergy between a wide range of cell types. Subtle differences between humans and mice (Chin *et al.*, 2017; Yanai *et al.*, 2017) combined with limited access to human fetal and embryonic tissue, has rendered our understanding of these processes in humans rudimentary. Importantly, environmentally triggered alterations in early developmental have been implicated in a range of immune mediated pathologies including inflammatory bowel diseases (IBDs) (Sonntag *et al.*, 2007; Cilieborg, Boye and Sangild, 2012; Dupaul-Chicoine, Dagenais and Saleh, 2013; Kraiczy *et al.*, 2016). Furthermore, a number of studies have reported a direct link between early fetal intestinal epithelial cell dynamics and IBDs (Kraiczy *et al.*, 2016; Yui *et al.*, 2018), suggesting that fetal-like transcriptional programmes may re-appear in the intestinal epithelium of patients diagnosed with IBD. Hence, deciphering physiological intestinal development is a critical first step towards either preventing such conditions or developing better treatments.

The human intestinal tract develops from the endodermal germ cell layer of the embryo, beginning with the formation of a simple tube at 3-4 weeks post-conception (PCW). Prior to villus formation, the intestinal epithelium, forming the innermost lining of the gut tube, is pseudostratified and has been shown to be globally proliferative (Grosse *et al.*, 2011). By the end of the first trimester (12 PCW), regionalisation of the intestinal tube has occurred and a crypt-villus axis has started to appear. Although the exact mechanisms of gut elongation and crypt-villus formation in humans remain unknown, key signalling events in mice include gradients of hedgehog (HH), WNT, FGF, and BMP (Korinek *et al.*, 1998; Madison *et al.*, 2005; Geske *et al.*, 2008; Shyer *et al.*, 2015). The exact composition of the epithelium, including transcriptional profiles of cells that give rise to the fully differentiated single cell layer, have yet to be described in humans.

In the adult intestine, *LGR5* is a marker of intestinal stem cells that reside at the bottom of intestinal crypts and give rise to all epithelial cell subsets (Barker *et al.*, 2007). The exact timepoint at which these cells appear in humans, and from which cells they are derived, remains to be determined. The ability to generate self-organising, intestinal epithelial organoids from fetal gut epithelium as early as 8-10 PCW implies the presence of epithelial cells that are capable of giving rise to such three-dimensional structures (Fordham *et al.*, 2013). Indeed, the use of organoid models as tools to investigate early fetal intestinal development has been demonstrated previously (Kraiczy *et al.*, 2019). Nevertheless, the cross-talk between epithelial cell subsets and other mucosal cell types as well as cell lineage trajectories remains unknown. Recent advances in the field of single-cell sequencing (scRNAseq) have opened novel opportunities to shed further light on these complex processes. Recent studies have used scRNAseq to interrogate intestinal regional specification in mice and humans (Gao *et al.*, 2018; Nowotschin *et al.*, 2019). However, human crypt-villi formation and epithelial dynamics have not yet been explored.

In this study, we performed single-cell transcriptional profiling of human embryonic and fetal gut samples obtained from 9 human embryos spanning ages 6-10 PCW. Additionally, we profile mucosal biopsies from the small bowel of healthy children aged 4-12 years as well as a group of children newly diagnosed with Crohn’s disease (CD), a common form of IBD. In total, we generate single cell transcriptomes of ∼90,000 primary human intestinal cells providing a rich resource and detailed roadmap. Using this data as well as scRNAseq profiles of human fetal gut derived organoids we describe embryonic and fetal epithelium composition, trace their differentiation dynamics and signalling partners, and provide links to regenerating Crohn’s disease epithelium.

## Results

### Single cell map of the human embryonic, fetal and paediatric gut

Human embryos with a post-conceptional age ranging from 6 to 10 weeks were dissected to remove the intestinal tube, which was divided into proximal small bowel (duodenum and jejunum), distal small bowel (ileum) and large bowel (colon). Additionally, we obtained small bowel (i.e. terminal ileum; TI) mucosal biopsies from healthy children aged 4 to 12 years undergoing routine endoscopic investigation to rule out gut diseases (Figure 1A and Supplementary Table 1). Tissue samples were treated using an enzymatic dissociation protocol and single cell suspensions were then processed using the 10x Genomics Chromium workflow (Materials and Methods). In a subset of samples, the intestinal epithelial cell fraction was enriched by performing magnetic bead sorting for EPCAM (Figure 1B and Supplementary Table 1). In total 62,854 fetal (n = 34) and 11,302 paediatric terminal ileal (*n* = 8) cells passed quality control and doublet exclusion (Extended Fig. 1, Supplementary Table 1).

**Figure 1:**
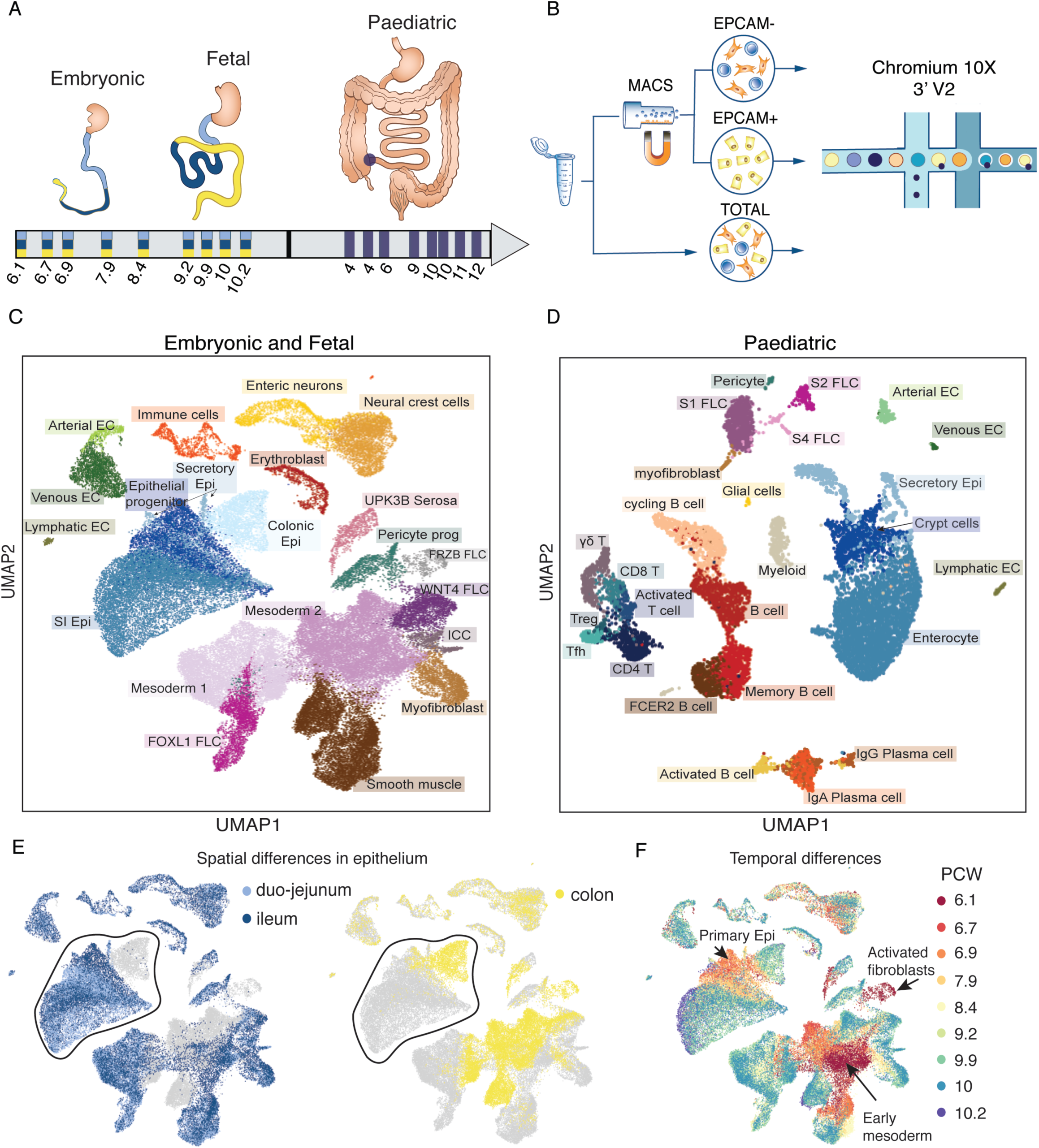
Single-cell profiling of embryonic, fetal and paediatric gut. A) Schematic illustration of experimental design, highlighting anatomical location and age of embryonic, fetal and paediatric samples collected. The blue circle marks the approximate biopsy location (i.e. terminal ileum). B) Tissue dissociation and single-cell sequencing strategy. C) and D) UMAP (uniform manifold approximation and projection) projects for embryonic/fetal (n = 9 donors) and childhood/adolescence (n = 8 donors) scRNAseq samples, respectively. UMAP plots coloured by E) gut region of the embryonic cells, F) post conception week of embryonic and fetal cells as in (C). Circled populations in E are epithelial cells. EC = endothelial cell, FLC = fibroblast cell, Epi = epithelium, SI = small intestinal, ICC = interstitial cells of Cajal.

Fetal and paediatric datasets were processed individually to identify the cell types present. Clustering and cell type-specific marker gene expression revealed 7 major cell types in fetal samples, including immune, erythroblast, endothelial, neural crest, smooth muscle, mesenchymal and epithelial cell populations (Extended Fig. 2A-D). Assessment of cellular subsets and their expression markers allowed us to further subdivide cell types as outlined in Figure 1C and D. All major cell types were also identified in paediatric TI biopsy samples with the exception of enteric neurons, smooth muscle and serosa cells (Figure 1D), whose exclusion would be expected given that the depth of forceps biopsy is restricted to mucosa.

**Figure 2:**
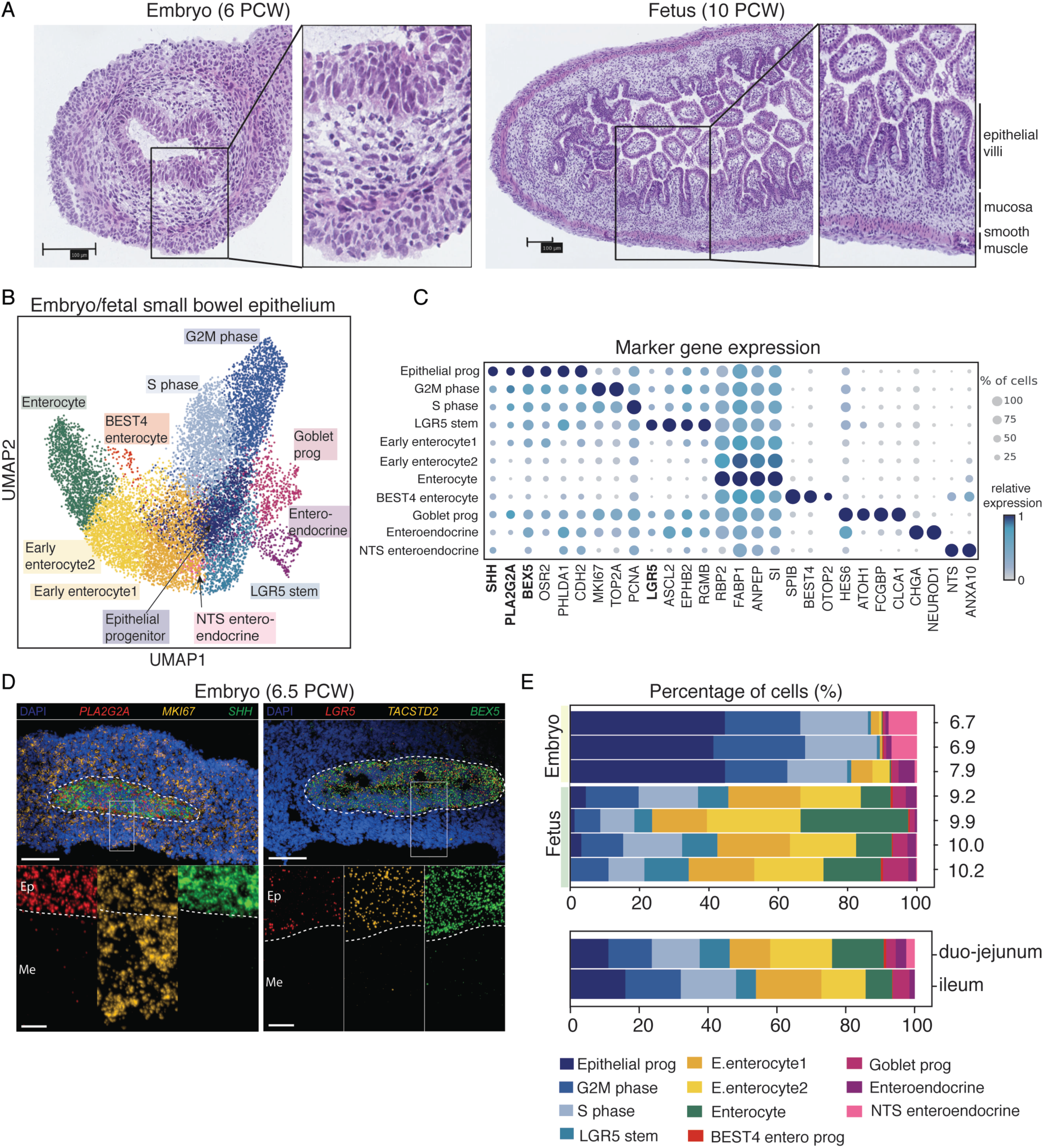
Changes in epithelial cell composition during crypt-villus formation in humans. A) Representative hematoxylin and eosin staining of embryonic and fetal ileum at 6 and 10 PCW (n = 3 donors). B) Sub-clustered epithelial cells from duo-jejunum and ileum colored by cell-type annotation. C) Dotplot with marker genes used to annotate fetal epithelial cell subtypes. D) Confocal microscopy images with single-molecule RNAscope staining of key marker genes expressed at pseudostratified epithelium stage. E) Changes in epithelial cell type abundance (% of cells) at different developmental time points and in two small bowel regions. Colors match the cell type annotation in B).

Comparing cellular composition across the three developmental stages (embryonic, fetal and paediatric), we observed striking differences. For example, the mesenchymal cell compartment was greatly expanded in proportion as well as diversity in embryonic and fetal samples (Figure 1C and D). Conversely, paediatric samples were dominated by immune cells including follicular/memory B cells, T cells, plasma and myeloid cells. Embryonic and fetal samples contained a comparatively small proportion of immune cells including macrophages, dendritic cells, monocytes as well as T and B cells (Data not shown).

As shown in Figure 1E, the spatio-temporal distribution of individual cell clusters demonstrated a significant separation of fetal epithelial cell clusters according to gut region as well as developmental time point (Figure 1E, Extended Fig. 3). Temporal separation in paediatric samples was less pronounced (Extended Fig. 1D). These differences highlight major changes that the intestinal epithelium undergoes during the captured developmental time periods. Hence, we next aimed to further elucidate underlying mechanisms and pathways.

**Figure 3.**
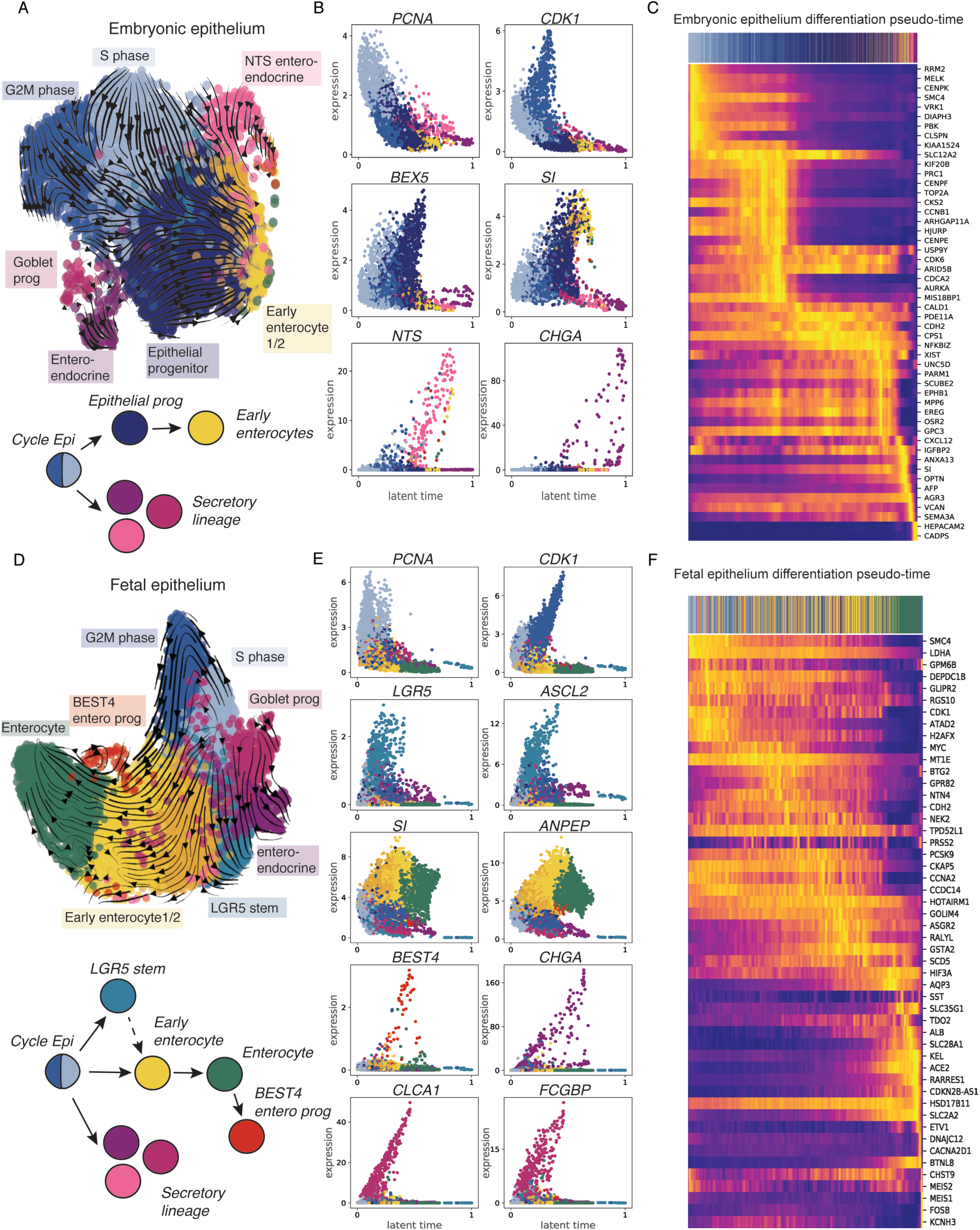
RNA velocity analysis of embryonic and fetal intestinal epithelium. A) and D) RNA velocities projected on the UMAP embedding of intestinal epithelium at embryonic and fetal stages, respectively. Schematics show summary of cell type differentiation dynamics. B) and E) Marker gene expression along the pseudotime (latent time). Cells are colored by the cell type as in UMAP. C) and F) Top 50 genes differentially expressed throughout pseudotime in embryonic and fetal epithelium, respectively. Top bars in (C) and (F) represent cell annotation.

### Intestinal epithelial cell type changes during human crypt-villus formation

Approximately midway through the first trimester, the human intestine is lined by a thick, pseudostratified epithelium that largely fills the intestinal lumen (Figure 2A). Only 3-4 weeks later a single-cell layer starts to appear, and by 10 PCW a primitive villus structure can be observed (Figure 2A). In order to examine transcriptional changes that occur during this phase of development and identify underlying regulatory networks, we sub-clustered fetal small bowel (duo-jejunum and ileum) epithelial cells based on the expression of *EPCAM*. As shown in Figure 2B-C, the pseudostratified epithelium primarily contains cells that are unique to developing epithelium and are defined by high expression of *SHH, PLA2G2A, BEX5*, and *CDH2*, as well as a high proportion of rapidly cycling epithelium (G2M and S phase). We further confirmed *in situ* expression of selected marker genes of the pseudostratified epithelium using multiplexed RNAscope (Figure 2D).

Despite the presence of cell cycle genes, *LGR5* expression was found to be low or absent in this epithelial cell state. Strikingly, this population of epithelial progenitor cells rapidly decreased at 10 PCW (Figure 2E), coinciding with early villous formation and the appearance of *LGR5+* stem cells, immature and maturing enterocytes, Goblet and enteroendocrine progenitor cells (Figure 2B and C). Although no Paneth cells were found during the first trimester of development, we observed a population of *BEST4*/*OTOP2*+ enterocytes, both findings confirming previous reports (Mallow *et al.*, 1996; Parikh *et al.*, 2019; Smillie *et al.*, 2019). Interestingly, we observed distinct differences in epithelial cell proportions by gut location with a higher abundance of progenitor cells present in distal compared to proximal small bowel samples (Figure 2E), suggesting that gut development follows a proximal to distal temporal gradient.

In mice, Trop2-expressing intestinal epithelial cells have been shown to be precursors to classical *Lgr5*+ stem cells (Mustata *et al.*, 2013). We applied the scVelo algorithm to fetal epithelial cells of the small bowel (duo-jejunum and ileum) to better understand the epithelial cell differentiation dynamics during the transition from embryonic (6-8 PCW) to fetal (9-10 PCW) epithelium (Figure 3A-F). At the embryonic stage, rapidly-cycling cells form the start-point of the trajectory and give rise to *BEX5*+ progenitor cells as well as a proportion of differentiated cells such as enteroendocrine and Goblet cell progenitors (Figure 3A and B). This suggests that pseudostratified epithelium is capable of expanding and self-renewing as well as giving rise to differentiated cell lineages, even in the absence of *LGR5+* cells, supporting a recent report in mice (Guiu *et al.*, 2019). Furthermore, the dynamic gene expression profile during early differentiation of the pseudostratified epithelium (Figure 3C), reveals rapid loss of the aforementioned progenitor cell markers during differentiation. At the fetal stage, cycling cells at the start-point of the trajectory differentiate into *LGR5+* stem cells and subsequently into its differentiated progeny (Figure 3D-F). This finding indicates that the cycling epithelium undergoes transcriptional transition from the pseudostratified epithelium into *LGR5+* stem cells and therefore may act as both a primitive stem cell of the early gut, and also as a progenitor to *LGR5+* stem cells later in development.

In summary, results of RNA velocity analyses reveal complex cell dynamics in the embryonic and fetal epithelium and further support the conclusion that a large proportion of pseudostratified intestinal epithelium at 6 PCW represents a rapidly cycling epithelial progenitor cell.

### Cell-cell cross-talk that supports crypt-villus formation in humans

Development of the human intestinal epithelium critically relies on the cross-talk with non-epithelial cell subsets. The most extensively studied example of such interactions in mammals is between epithelial and mesenchymal cells that regularly cluster and restrict epithelial cell proliferation at the top of the forming villi (Karlsson *et al.*, 2000; Walton *et al.*, 2012; Walton, Freddo, *et al.*, 2016). We next aimed to investigate the cross-talk between the intestinal epithelium and surrounding cell-types with a focus on mesenchymal cells. Using the previously described CellPhoneDB algorithm (Efremova *et al.*, unpublished Vento-Tormo *et al.*, 2018), we found that the three mesenchymal populations displaying the highest number of cell-type specific interactions with epithelial cell types were *FOXL1+* and *WNT4+* fibroblasts and *UPK3B*+ serosal cells (Figure 4A). In our dataset, *FOXL1*+ fibroblasts expressed developmentally related genes such as *DLL1* (Notch signaling ligand) and platelet derived growth factor receptor (*PDGFR*) (Karlsson *et al.*, 2000; Kurahashi *et al.*, 2008) as well as adult colonic mucosal S1 fibroblast markers including *F3* and *ADAMDEC1* (Kinchen *et al.*, 2018). At 6 PCW, *FOXL1* expressing fibroblasts were located in close proximity to the intestinal epithelium, as confirmed *in situ* using RNAscope (Figure 4B). While this population expressed some WNT2B signals, another population high in WNT was mesothelial serosa cells. This population showed expression of both epithelial keratin (*KRT8*) and mesenchymal collagen (*COL1A1*) genes and was uniquely marked by *UPK3B* expression (Extended Fig. 4). We confirmed their spatial localisation at the outermost layer of the intestinal tube using RNAscope (Figure 4B).

**Figure 4:**
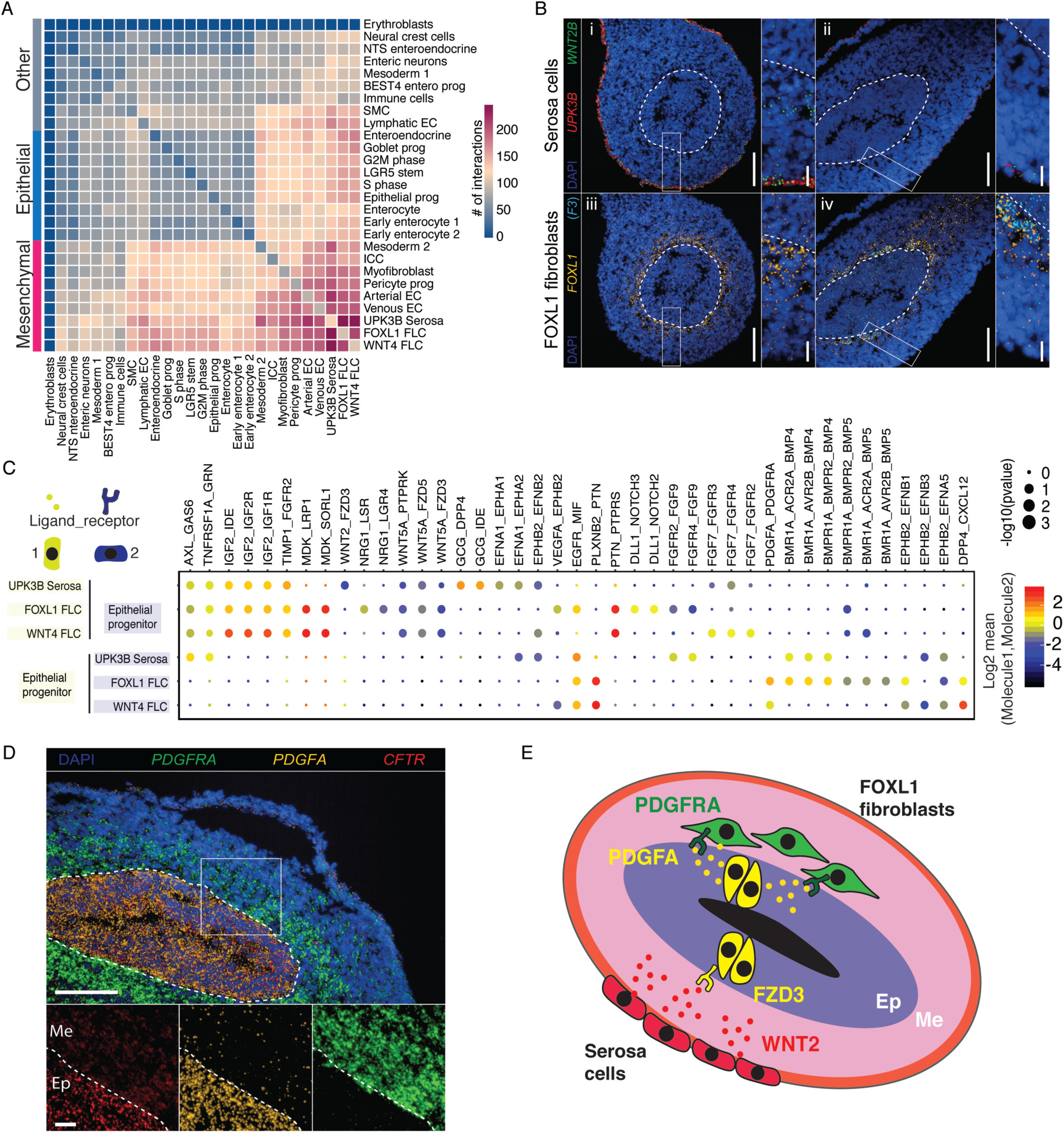
Cell-cell interactions that support transition from embryonic to fetal epithelium in humans. A) Heatmap showing a number of significant (or cell-type specific, p-value ≤0.05) interactions captured between cells found in developing human intestine as identified using CellphoneDB. B) RNAscope images of markers expressed by *FOXL1*+ fibroblasts (*F3* and *FOXL1*) and *UPK3B*+ serosa cells (*UPK3B* and *WNT2B*) in developing proximal intestinal tissues. C) Dotplot of ligand-receptor interactions between epithelial progenitor cells and three mesenchymal populations predicted using CellPhoneDB analysis. Point size indicates permutation *p* value and color indicates the scaled mean expression level of ligand and receptor (Mol1/2). D) RNAscope images of *PDGFRA* (mesenchymal receptor) and *PDGFA* (epithelial ligand) interacting pair predicted by CellphoneDB. Epithelium distinguished by *CFTR* expression. E) Schematic showing predicted ligand-receptor interactions between epithelial (yellow), mesenchymal (green) and serosa (red) cells. FLC = fibroblasts.

The most specific interactions between *FOXL1+* fibroblasts and the epithelial progenitor cells were via bone morphogenetic protein (BMP) (e.g. *BMPR1A-BMPR2-BMP5*), Wnt (e.g. *WNT5A-FZD5*/3), FGF (*FGF9-FGFR2/4*) and Notch (e.g. *DLL1-NOTCH 2/3*) signalling (Figure 4C). These findings are in keeping with previous reports derived from mouse models of gut development highlighting these signalling pathways as critical gatekeepers of epithelial cell transition and villus formation (Geske *et al.*, 2008; Walton, Whidden, *et al.*, 2016; Kaestner, 2019). Other specific ligands secreted by *FOXL1+* fibroblast included *NRG1, PTN* and *VEGFA*. In addition, we found that serosa cells uniquely signaled to epithelial progenitor cells via glucagon (GCG) and WNT2 ligands, while *WNT4+* fibroblasts expressed *FGF7* and *CXCL12* ligands (Geske *et al.*, 2008). We also identified growth factors, such as *IGF2, PTN, MDK*, common to these mesenchymal subsets and received by epithelial progenitor cells (Figure 4C).

Lastly, platelet-derived growth factor receptor alpha (PDGFRA) and its ligand PDGFA have been implicated in early gut development and specifically villus morphogenesis in mice (Karlsson *et al.*, 2000). Our single-cell data and *in situ* imaging suggests that similar mechanisms are operative in humans, such that the early epithelial progenitor cells initiate villus morphogenesis by secreting PDGF ligands and thereby signalling to the underlying mesenchymal cells (Figure 4D-E).

Together, the data show interactions relevant to human crypt-villus formation and point towards existing and novel signalling pathways implicated in early human fetal gut development.

### Fetal organoids show *in vitro* maturation recapitulating *in vivo* epithelial transition

The ability of intestinal epithelial stem cells to give rise to all cell subsets has led to the development of the intestinal organoid culture model (Sato et al., 2011). Such organoids can also be generated from human fetal gut, providing the opportunity to investigate epithelial cell-intrinsic and extrinsic developmental mechanisms (Fordham et al., 2017, Kraiczy et al., 2018). Here we applied scRNAseq to human fetal gut organoids (Material and Methods, Supplementary Table 1) and performed *in silico* analyses by classifying epithelial cells using transcriptional profiles derived from primary tissue.

In the adult small bowel mucosa, Paneth cells have been found to express the Wnt-agonist WNT3A, thereby providing a critical signal to the stem cell niche (Sato, van Es, *et al.*, 2011). Interestingly, we were unable to identify any *WNT3A* expressing cell types in our fetal gut scRNAseq datasets. However, given that WNT3A forms a key ingredient of previously reported human adult and fetal mucosa-derived intestinal epithelial organoid cultures (Sato, Stange, *et al.*, 2011; Fordham *et al.*, 2013; Kraiczy *et al.*, 2019) we aimed at assessing its requirement and impact on fetal gut organoids. Intestinal epithelial organoid (IEO) cultures were generated from the proximal small bowel and cultured either in the presence or absence of WNT3A conditioned medium. Inclusion of WNT3A led to the presence of budding, crypt structures while organoids lacking WNT3A appeared more spheroid-like (Figure 5A). Interestingly, single-cell transcriptional profiling of these cultures at an early passage (i.e. passage 2 – approximately 2-3 weeks in culture) revealed the presence of intestinal epithelial cells as well as a small fraction of mesenchymal cell populations as indicated by clear clustering of cells and expression of key epithelial and mesenchymal cell markers, respectively (Figure 5B and C, Extended Fig. 5A,B). Organoids were viable for several weeks even if cultured in the absence of WNT3A, and showed evidence of active cell cycling (Figure 5B, right panel). However, WNT3A was required for long term culture as WNT3A negative organoids showed reduced viability and could not be cultured beyond 6 weeks. Importantly, observed phenotypic differences were matched by dramatic transcriptional changes leading to a clear separation of cells according to culture conditions (Figure 5B). Removing or reducing WNT3A in adult mucosa-derived IEOs has been shown to induce differentiation of epithelial cells and a reduced expression of LGR5 (Kraiczy *et al.*, 2019). In contrast, when assessing epithelial cell identity and composition of fetal organoids using a logistic regression model based on our primary fetal scRNAseq profiles, organoids cultured in the presence of WNT3A were found to contain a greater proportion of differentiated cell types including enterocytes and enteroendocrine cells compared to those cultured in its absence (Figure 5D, left panel). Similarly, the predicted composition of mesenchymal cell fraction varied substantially between conditions with the majority of cells being identified as FOXL1+ fibroblasts in both conditions (Figure 5D, right panel).

**Figure 5:**
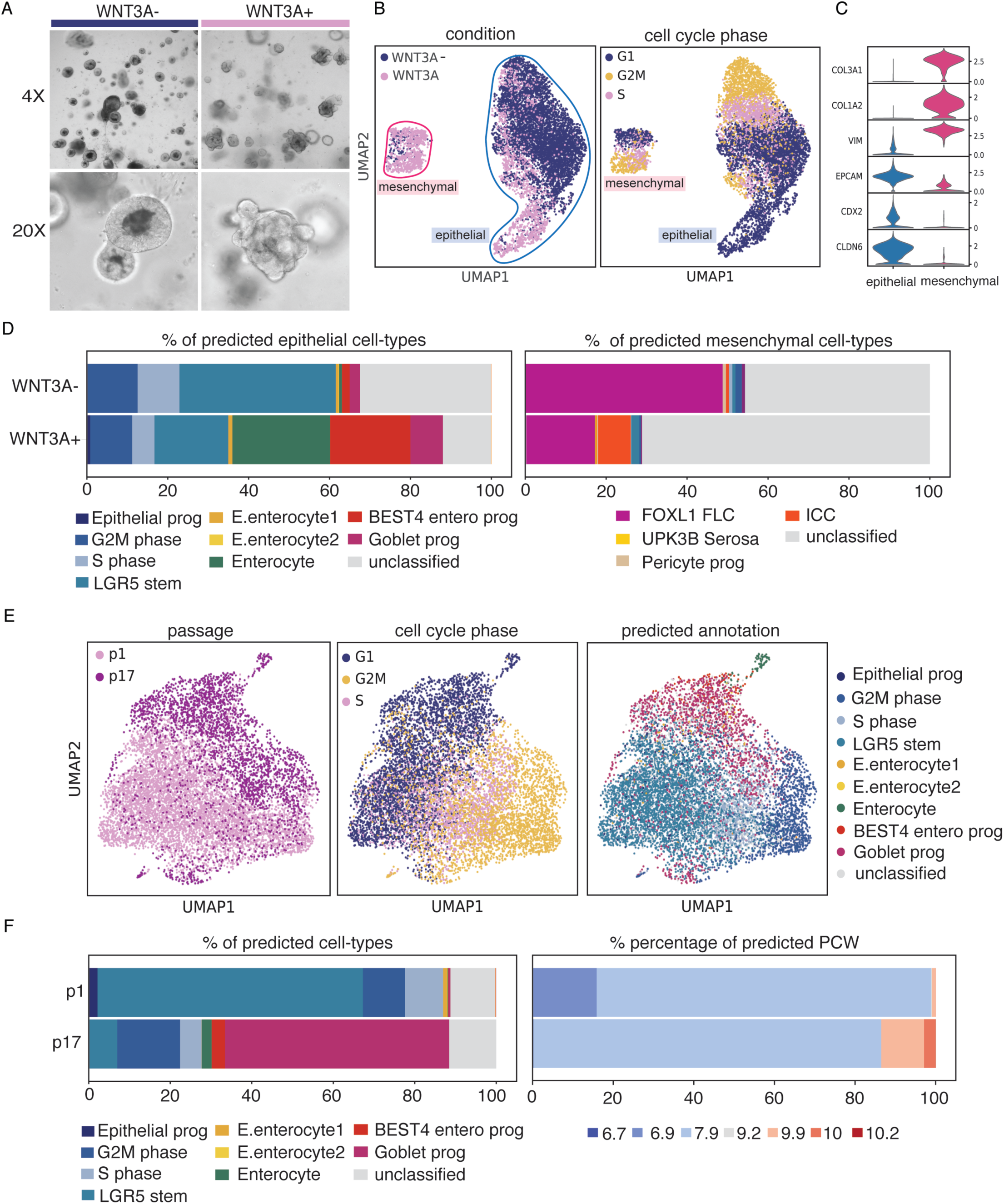
Fetal intestinal organoids mature in culture. A) Brightfield images of fetal organoids grown without (WNT3A-) or with (WNT3A+) conditional medium at passage 2. B) UMAP plot of single cells from fetal organoids grown with or without WNT3A. Cells are colored by either condition or cell cycle phase. C) Violin plots with mesenchymal and epithelial gene expression in two clusters (circled blue and pink in B). D) Abundance of cell types (% of cells) in organoids, as predicted by the logistic regression classifier trained on primary fetal cells. E) UMAP plots with single cells of fetal organoids grown for 1 and 17 passages and colored by passage, cell cycle phase or predicted annotation. Unclassified label indicates cells with prediction probability lower than 50%. F) Predicted cell type abundance or PCW age of fetal organoids grown for 1 or 17 passages as predicted using logistic classifier.

Previous work suggests that human fetal gut organoids undergo a degree of *in vitro* maturation in culture (Kraiczy *et al.*, 2019). In order to examine this phenomenon further, we generated organoids from embryonic gut samples aged 6 PCW and kept them in complete culture medium (containing WNT3A) over 5 months (17 passages). Single-cell profiling was applied to cultures once they were first established (after one week, passage 1) and following 5 months in culture. UMAP clustering revealed separation according to duration in culture, suggesting that significant transcriptional changes occur in culture (Figure 5E). Major differences were also observed with regards to the predicted cell composition, such that older cultures contained a higher proportion of differentiated cell types including enterocytes and goblet cell progenitors (Figure 5E, right panel, and 5F, left panel, Extended Fig. 5C). Importantly, estimated gestational age of organoid cultures was found to be higher at late passages providing further evidence for a degree of maturation occurring *in vitro* (Figure 5F, right panel).

In summary, our findings reveal effects of Wnt signalling and specifically WNT3A on human fetal epithelial organoid developmental stage and cell diversity.

### Reactivation of fetal transcriptional pathways in the inflamed epithelium of CD patients

Alterations in the composition, function and cell dynamics of the intestinal epithelium are thought to play a critical role in the pathogenesis of IBDs, including Crohn’s disease (CD). Moreover, a link has been established between early fetal development and the intestinal epithelium of IBD patients by demonstrating partial reprogramming of the regenerating epithelium into a primitive state (Yui *et al.*, 2018). In order to investigate these findings further we performed scRNAseq small bowel (terminal ileum) biopsies obtained from children newly diagnosed with CD (n = 7, Extended Figures 1B, 1D, and 6) and analysed the epithelial cell fraction. Comparing epithelial cell composition between CD patients and age matched non-IBD (healthy) control patients, we observed significant differences including an increase in transit amplifying (TA), Goblet and Tuft cells while the proportion of fully differentiated enterocytes was significantly reduced in the CD epithelium (Figure 6A-C, Extended Fig. 7C-D). Next, we interrogated the cross-talk between fibroblasts and the affected intestinal epithelial subtypes in the context of CD. We identified a number of cell-receptor interactions that were present only in CD patients (Figure 6D). For example, S4 fibroblasts were expanded in IBD (Extended Fig. 7C) and found to uniquely signal to CD TA cells via WNT2 ligands that are received via FZD3 receptor (Figure 6D). In addition, we observed chemokine ligands such as CXCL2, CXCL10 and CCL2 were expressed by the S4 fibroblast population and received by intestinal epithelial cells. Together, these findings highlight the complex cross-talk between the epithelium and surrounding cells which is likely to contribute towards chronic relapsing mucosal inflammation observed in childhood-onset CD.

**Figure 6:**
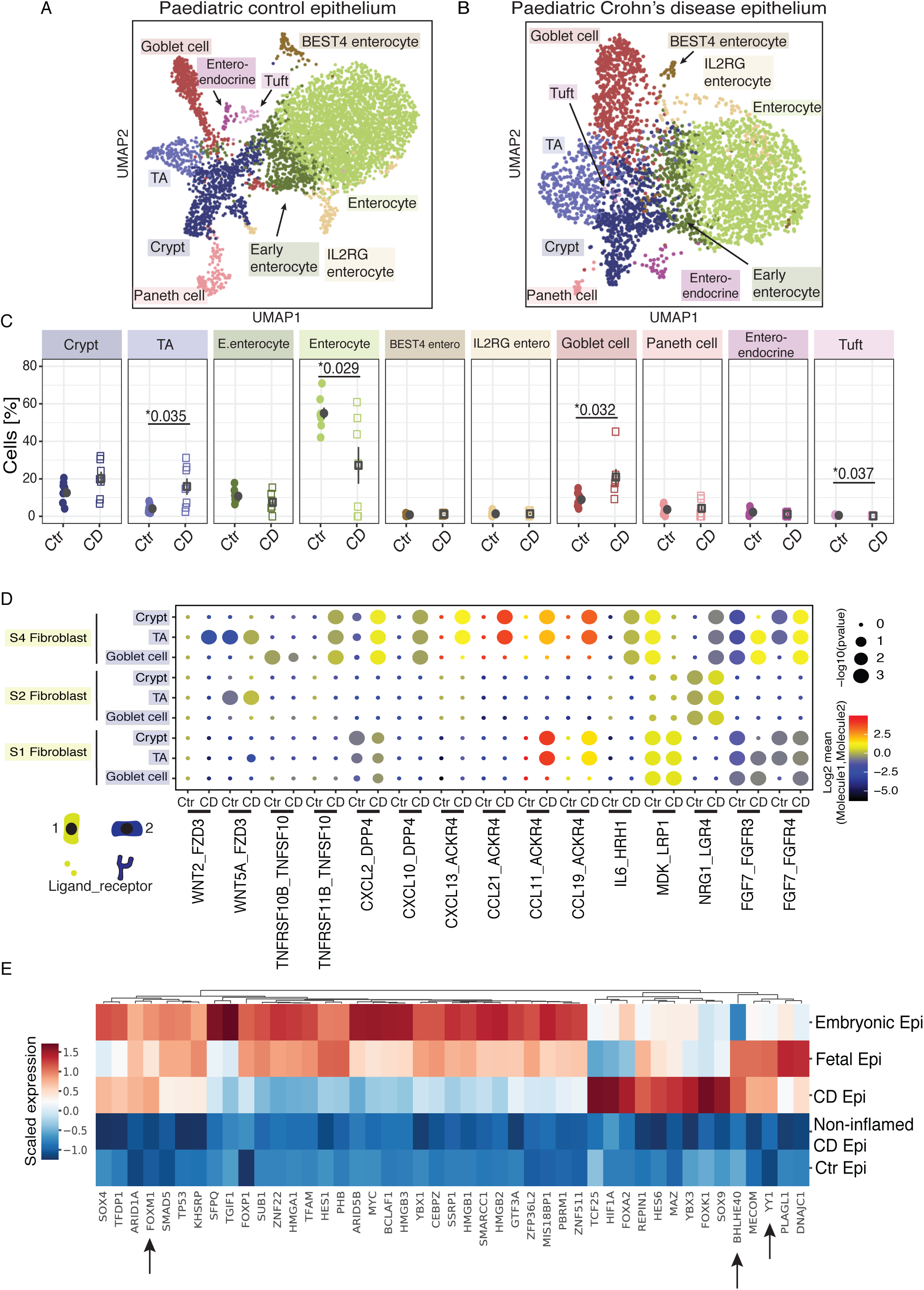
Epithelial cell dynamics in Crohn’s disease patients show transcriptional similarities to developing epithelium. A) and B) UMAP plots of epithelial cell subtypes in healthy children (n=8) and patients with Crohn’s disease (CD) (n=7), respectively. C) Epithelial cell type changes in paediatric health and CD. Transit amplifying (TA), enterocytes, goblet cells and tuft cell proportions were significantly changed between control and CD patients (p-values indicated, t-test). d) Dot plot with ligand-receptor interactions between S4 FLC, S2 FLC and S1 FLC and selected epithelial cell types. Point size indicates permutation *p* value (CellPhoneDB). Color indicates the scaled mean expression level of ligand and receptor. E) Heatmap shows relative mean expression of transcription factors, which were identified to be differentially expressed in CD epithelium, across epithelium from five groups. “Non-inflamed CD” were a group of patients with minimal epithelial composition changes as in (Extended Fig. 7D, arrows). Arrows mark genes relevant in CD and development.

Finally, we aimed to identify the transcription factors that are actively expressed in the CD but not healthy paediatric epithelium that are shared with embryonic or fetal epithelium (Materials and Methods). Indeed, as shown in Figure 6E, we identified a number of such genes, many of which have been previously reported as being expressed in the fetal intestinal epithelium and linked to IBD pathogenesis. Examples include genes encoding the B lymphocyte-induced maturation protein-1 (Blimp1; encoded by the gene *PRDM1*) (Harper *et al.*, 2011; Muncan *et al.*, 2011; Ellinghaus *et al.*, 2013), Wnt signalling transcription factor TCF4 (Barker *et al.*, 1999; Wehkamp *et al.*, 2007) as well as C-MYC, a transcription factor controlling the differentiation of progenitor cell types (Bettess *et al.*, 2005; Matthews *et al.*, 2019). Furthermore, our analyses also identified several novel transcription factors, expression of which becomes reactivated in the inflamed, regenerative CD epithelium.

Taken together, our results confirm previous reports of altered intestinal epithelial cell dynamics and identifies several known and novel disease-associated cell-to-cell interactions in childhood-onset CD. Importantly, we provide supportive evidence for the partial reactivation of primitive transcriptional pathways and demonstrate altered epithelial cell composition particularly in the inflamed, regenerating CD epithelium.

## Discussion

Recent single-cell studies have been instrumental in interrogating critical events during intestinal development in humans, including intestinal regionalisation (Gao *et al.*, 2018) and immune cell landscape development (Li *et al.*, 2019; Schreurs *et al.*, 2019). Here, we zoom-in on the first trimester development of the human intestine with the focus on epithelial cell type composition and dynamics during crypt-villus formation. In doing so, we describe transcriptional profile of the embryonic progenitor cell and its differentiation dynamics both *in vivo* and *in vitro*, as well as cell-cell interactions active during intestinal development and relevant during crypt-villus formation. In addition, we compare fetal and paediatric CD epithelium and identify a number of transcription factors expressed during development and re-activated during epithelial remodelling in CD. We provide single-cell data on the website **www.gutcellatlas.org** as a resource for the scientific community to address a range of research questions in the context of intestinal development, health and disease

### Single cell transcriptomic profile of epithelial progenitor cell

Previous studies have reported the presence of proliferative, immature progenitor epithelial cells in the human fetal intestine at around 10 PCW (Fordham *et al.*, 2013, Guiu *et al.*, 2019). We demonstrate that this cell population forms the vast majority of the pseudostratified intestinal epithelium in the human embryonic stage (6-8 PCW). Furthermore, this population was marked by strong expression of CDH2, BEX5, SSH and PLA2G2A, all of which have been previously linked to stem cell potential. For example, CDH2 was linked to regulation of cell fate decision in the mesodermal lineage (Alimperti and Andreadis, 2015), and SHH to initiation of villus formation in the developing mouse intestine (Shyer *et al.*, 2015), while BEX family genes were found to be expressed in tissue stem/progenitor cells (Ito *et al.*, 2014). Although this cell type was found to be rapidly cycling, their transcriptional profile differed distinctly from LGR5+ epithelial cells, which were found to be present in small numbers at 6 PCW.

Coinciding with major anatomical changes, we observed a dramatic change in cellular composition of epithelium during development at 7-9 PCW. In contrast to embryonic epithelium, the single cell layer lining primitive fetal villi contained a higher proportion of LGR5+ cells as well as early forms of differentiated epithelial cell subsets. This suggests that epithelial progenitor cells transition to LGR5 stem cells as has been proposed previously (Fordham *et al.*, 2013; Guiu *et al.*, 2019). Pseudotime analysis suggested that these primitive epithelial cells can give rise to differentiated cell subsets (such as enterocytes, Goblet and enteroendocrine cells) either directly or via LGR5+ intermediate cells. These findings are consistent with recent reports that demonstrate that both villus and inter-villus epithelial cells are equipotent and have the same regenerative potential (Guiu *et al.*, 2019), therefore suggesting that properties of LGR5 stem are induced by their surroundings.

Using *in silico* ligand receptor analyses, we further identified *FOXL1* and *WNT4A* expressing fibroblast as well as serosa cells populations as key cell types providing key stem cell niche ligands including *WNT5A, WNT2, NRG1* and *IGF2*. In mice, *Foxl1*+ subepithelial telocytes provide Wnt signals to the crypt epithelial cells in adults (Shoshkes-Carmel *et al.*, 2018) and during development (Kaestner *et al.*, 1996, 1997; Fukamachi *et al.*, 2001). Our imaging data suggests that this cell type is already present at the embryonic stage in the absence of any visible villi and hence, suggests that these cells may provide a critical first signal towards villi formation. We also show that prior to the development of *FOXL1*+ telocytes, the major source of WNT2B is serosa cells that surround the outermost layer of the gut. Whether the signals are able to travel to epithelium or affect adjacent differentiating mesenchymal populations is unknown.

### Human fetal gut derived organoids as models to investigate epithelial cell development

Intestinal epithelial organoids have been generated from human fetal gut and shown to undergo a degree of *in vitro* development highlighting their use as powerful experimental tools (Fordham *et al.*, 2013; Guiu *et al.*, 2019; Kraiczy *et al.*, 2019). Here we combine generation of fetal organoids with single-cell profiling to interrogate fetal culture composition. Our findings indicate that whilst not required for the establishment and short-term culture, WNT3A is essential for long-term propagation of fetal organoids and was found to be associated with a higher proportion of differentiated cell subsets. This parallels studies in mice, where embryonic progenitors were able to proliferate independently of Wnt prior to villi formation but not after (Chin *et al.*, 2016). These findings also point to differences between adult and fetal gut epithelium as the generation of adult mucosa derived intestinal organoids critically relies on the presence of WNT3A while its withdrawal leads to increased differentiation into epithelial cell subsets and reduced expression of LGR5 (Fordham *et al.*, 2013; Kraiczy *et al.*, 2019). Furthermore, organoids kept in culture for several months were found to contain an increased proportion of differentiated cell-subsets as well as an increased number of LGR5+ cells. This suggests that current intestinal culture conditions select for highly proliferating cells. Finally, our *in vitro* studies further illustrate the utility of single cell transcriptomic maps as a critical reference to validate and interpret fetal organoid culture work.

### Linking fetal gut epithelial development to childhood-onset CD

A developmental origin of disease pathogenesis has been proposed for many complex, multifactorial conditions. IBDs, and particularly CD, are thought to be caused by a complex interplay between the environment and genetic predisposition leading to an irreversibly altered immune response. Recent studies have reported expansion of colonic mesenchymal subset in adult ulcerative colitis and associated this with resistance to anti-TNF treatment (Kinchen *et al.*, 2018; Smillie *et al.*, 2019). We also observe an expansion of this mesenchymal population in childhood Crohn’s disease, suggesting similarities between adult- and paediatric-onset IBD as well as their subtypes. In addition, comparison between CD and healthy epithelium suggests that CD epithelium is rapidly cycling and poised for Goblet cell differentiation. Furthermore, we describe ligand and receptor pairs that uniquely signal between affected epithelial subsets and expanded fibroblast populations, including WNT2 ligands received by transit amplifying cells. This provides a possible mechanism to sustain intestinal regeneration in disease.

Previous studies in mice have linked epithelial cell properties in the inflamed gut to the physiological status observed in early fetal development (Yui *et al.*, 2018; Guiu *et al.*, 2019). Here, we provide evidence in humans that regenerating CD epithelium shares transcription factor programs otherwise present only in fetal epithelium. Amongst them is the zinc finger transcription factor Yin Yang (YY1), which has been shown to play a critical role in lung epithelial cell development and TGF-beta induced lung fibrosis (Boucherat *et al.*, 2015; Zhang *et al.*, 2019). Another example is the expression of basic helix–loop–helix 40 (BHLH40) in both fetal and CD epithelium. Interestingly, BHLH40 expression has been observed in a wide range of cells and tissues including T-cells, macrophages, dendritic cells and the gastric epithelium (Lin *et al.*, 2014; Teng *et al.*, 2020). BHLH40 was found to control cytokine production by T-cells thereby playing a critical role in the development of autoimmune neuroinflammation (Lin *et al.*, 2014; Yu *et al.*, 2018). Finally, our analyses confirm previous reports of Forkhead BoxM1 (FOXM1, also HFH-11) transcription factor being expressed in embryonic epithelial cell with its expressing becoming reactivated in adult cell types by proliferative signals or oxidative stress (Ye *et al.*, 1997).

In summary, we provide a detailed single cell map of the human gut during embryonic, fetal and paediatric health as well as inflammatory disease, and use the data to dissect epithelial cell dynamics during intestinal life.

## Acknowledgements

This work was financially supported by: a Horizon 2020 grant, (668294, ‘INtestinal Tissue ENgineering Solution for children with short bowel syndrome’, L.V.); an ERC Advanced Grant (New-Chol, L.V.); the Cambridge University Hospitals National Institute for Health Research Biomedical Research Centre (L.V.); a core support grant from the Wellcome Trust and MRC to the Wellcome Trust - Medical Research Council Cambridge Stem Cell Institute (L.V.); the Wellcome Trust (WT206194, S.A.T.);the European Research Council (646794, ThDefine, S.A.T.); an MRC New Investigator Research Grant (MR/T001917/1, M.Z.); and a project grant from the Great Ormond Street Hospital Children’s Charity, Sparks (V4519, M.Z.). We thank Dr Franco Torrente and Dr Camilla Salvestrini as well as Claire Gleams for recruiting paediatric patients and obtaining biopsy samples.

## Author Contribution

A.R., L.V., M.Z. and S.A.T initiated, designed and supervised the project. A.R., R.E. and K.N. performed tissue processing, organoid culture and scRNAseq experiments. R.E. analysed single-cell data and generated figures. T.G. contributed to data analysis. K.R.J., D.O. and T.G. supported analyses, critical discussion and interpretation of data. K.R., L.T. and O.A.B. performed tissue sectioning, staining and imaging. R.H. carried out paediatric patient recruitment, obtaining biopsy samples and documentation of clinical data. A.R., R.E. and M.Z. wrote the manuscript. All authors contributed to discussion and interpretation of results as well as editing of the manuscript.

## Material and methods

### Fetal and paediatric tissue sampling

First trimester human fetal tissue was collected from patients undergoing elective termination of pregnancy. Patients gave informed consent as part of the ethically approved research study (REC-96/085). Fetal age (post conception weeks, PCW) was estimated using the independent measurement of the crown rump length (CRL), using the formula PCW (days) = 0.9022 × CRL (mm) + 27.372.

Human intestinal mucosal biopsies were obtained from patients undergoing colonoscopy at Addenbrooke’s Hospital, Cambridge, UK. All patients gave informed consent for extra biopsy samples to be taken for research use when undergoing elective colonoscopy (REC 17/EE/0265). Patients were then included if, after macroscopic visualization and histological analysis, they were diagnosed as either having Crohn’s disease, or being without an inflammatory diagnosis (control).

### Fetal and paediatric tissue dissociation

Fetal intestinal samples were dissected into duodenum-jejunum (further referred to as the duo-jejunum), ileum and colon using anatomical landmarks, and processed to single-cell suspension in parallel. Both fetal samples and paediatric samples were processed using the same protocol. Briefly, paediatric biopsies or fetal tissue sections were immediately rinsed twice with Hank’s Balanced Salt Solution (HBSS) medium (Sigma-Aldrich) and digested in HBSS medium containing 1.07 Wünsch units/ml of Liberase DH (Roche) and 600 IU of Hyaluronidase (Calbiochem) on a shaking platform (750 rpm) at 37°C for up to 30 min. The tissue was gently homogenised using a P1000 pipette every 15 mins. A single-cell suspension was then passed through a 40 µm cell strainer to remove undigested tissue. Cells were spun down at 400 g at 4°C for 5 min and the pelleted cells were washed in DMEM/F12 three times using centrifugation.

Fetal cells were either loaded for scRNAseq directly following sample processing or subjected to EPCAM selection to enrich for epithelial cells. For enrichment, single cells were suspended in MACS modified solution (PBS with 0.5% BSA, 2 mM EDTA and 100 IU/mL DNaseII) and stained with EPCAM (CD326) magnetic microbeads (Miltenylbiotec) according to the manufacturer’s protocol. Enrichment was performed using an autoMACS Pro Separator. Either only EPCAM positive (PCW 6.7, 6.9, 10.2, 9.3) or both EPCAM positive and negative (PCW 9.9, 10.1, 10) fractions were processed using the 10x Genomics single-cell transcriptomics system. All paediatric single-cell suspensions were subjected to the MACS enrichment using the same protocol as described for fetal samples. Both fetal and paediatric single-cell suspensions were carried forward into single-cell sequencing only if the viability was >60% (Supplementary Table 1).

### Intestinal organoid culture

Fetal organoids were cultured according to a protocol by (Fordham *et al.*, 2013). During organoid culture, the media was replaced every 48-72 hours. For full media composition, see supplementary table 3. Organoids were passaged using mechanical disruption with a P1000 pipette and re-seeding in fresh growth-factor reduced Matrigel® (Corning). Processing for single-cell sequencing analysis was performed by removing the organoids from matrigel using incubation with Cell Recovery Solution at 4°C for 30 minutes, pelleting the cells, and re-suspending in TrypLE enzyme solution (Thermo Fisher) for incubation at 37°C for 15 mins. Cells were pelleted again and re-suspended in DMEM/F12. Cells from fetal ileum from embryos of 6 PCW were dissociated into single-cell suspension at passage 1 (∼1 week) or passage 17 (∼1 month) in culture and profiled using 10x Genomics single-cell transcriptomics. Brightfield images of organoids were taken using an EVOS Cell Imaging Systems microscope (Thermo Fisher).

### Tissue freezing, sectioning and RNAscope

Tissue was prepared, stained, and imaged largely as described previously (Bayraktar *et al.*, Nature Neuroscience, in press)). Fresh tissue samples were either embedded in OCT and frozen at −80°C on an isopentane-dry ice slurry, or fixed in 10% neutral-buffered formalin at 4°C for ∼24 hours, and then embedded and frozen as above. Cryosections were cut at a thickness of 10-16 µm using a Leica CM3050 S cryostat and placed onto SuperFrost®Plus™ slides (VWR). Prior to staining, tissue sections were post-fixed in 4% paraformaldehyde in PBS for 15 minutes at 4°C, then dehydrated through a series of 50%, 70%, 100%, and 100% ethanol, for 5 minutes each.

Tissue sections were then processed using a Leica BOND RX to automate staining with the RNAscope® Multiplex Fluorescent Reagent Kit v2 Assay and RNAscope® 4-plex Ancillary Kit for Multiplex Fluorescent Reagent Kit v2 (Advanced Cell Diagnostics, Bio-Techne), according to the manufacturers’ instructions. Automated processing of non-fixed sections included pre-treatment with Protease IV for 30 minutes, but no heat treatment; fixed frozen sections were subjected to heat-induced epitope retrieval at 95°C in buffer ER2, and digestion with Protease III for 15 minutes. Tyramide signal amplification with Opal 520, Opal 570, and Opal 650 (Akoya Biosciences) was used to develop three probe channels. The fourth was developed using TSA-biotin (TSA Plus Biotin Kit, Perkin Elmer) and streptavidin-conjugated Atto 425 (Sigma Aldrich).

Stained sections were imaged with a Perkin Elmer Opera® Phenix™ High-Content Screening System, in confocal mode with 1 µm z-step size, using 20× (NA 0.16, 0.299 µm/pixel) and 40× (NA 1.1, 0.149 µm/pixel) water-immersion objectives.

### Single-cell RNA-sequencing

Single-cell suspension for each primary intestinal or organoid sample was loaded onto a separate channel of a Chromium 10x Genomics single cell 3’v2 library chip as per manufacturer’s protocol (10x Genomics; PN-120233), aiming for a cell capture recovery of 3000-5000 cells. cDNA sequencing libraries were prepared according to the manufacturer’s protocol and sequenced on an Illumina Hi-seq 4000 (2×50bp paired-end reads).

### Processing FASTQ files and quality control

Raw sequence reads in FASTQ format from fetal, paediatric and organoid samples were processed and aligned to the GRCh38-1.2.0 human reference transcriptome using the Cellranger v2.1.1 pipeline (10x Genomics) with default parameters.

The resulting gene expression matrices integrated together using Scanpy package v1.4 (Wolf, Angerer and Theis, 2018). A total of 34 fetal sample count matrices were merged together. Separately, 15 gene expression matrices of healthy and Crohn’s disease paediatric biopsy samples were merged together for cell annotation and direct comparisons. Organoid gene expression matrices from the same experiment were also merged separately.

The pre-processing followed the guidelines provided by Scanpy V1.4 tutorial (Wolf, Angerer and Theis, 2018). In short, entries with fewer than 200 genes and greater than 9000 total genes were filtered to remove empty droplets and probable doublets, respectively. The distribution of Unique Molecular Identifiers (UMIs) and genes per cell were visualised using scanpy pl.scatter function (Supp. Figure 1 A-C). To account for differences in sequencing depth across samples, we normalised expression values for total UMIs per cell and log transformed the counts.

### Doublet removal

All 10x runs were processed using Scrublet doublet detection pipeline with threshold of 0.25-0.3 (Wolock, Lopez and Klein, 2019), and predicted doublets were excluded from the analysis. We further annotated doublets by sub-clustering the data and identifying clusters with gene expression of other clusters. In fetal samples the doublets were largely epithelial-mesenchymal and neuronal-mesenchymal. In paediatric samples, we found mostly T cell-enterocyte and Goblet cell-enterocyte doublets.

### Clustering, visualisation and cell annotation

For cell clustering we used highly variable genes selected using sc.pp.highly_variable_genes function with default parameters. In addition, cell-cycle signatures were determined using cycle stage marker genes imported from human_cycle_markers.rds, scran package (Scialdone *et al.*, 2015) and removed from highly variable genes of the full dataset. Similarly, ribosomal protein genes were removed from the highly variable genes as they contributed to the highest variability in the F6.1 sample. In addition, UMI counts, percentage of mitochondrial genes were considered to be the source of unwanted variability and were regressed using Scanpy regress_out function (Wolf, Angerer and Theis, 2018).

To remove variation of each 10X Genomics run and maintain the development related biological variation, we used batch balanced k nearest neighbour (BBKNN) method (Polański *et al.*, 2019) on 40 principal components and trim parameter set to 20. Dimensionality reduction was performed on remaining highly variable genes and cells were visualised using Uniform Manifold Approximation and Projection (UMAP) plots (Becht *et al.*, 2018). We then used Scanpy implementation of Leiden algorithm for unsupervised clustering of the data (Traag, Waltman and van Eck, 2019). Clusters were annotated using markers genes found in the literature in combination with differentially expressed genes (Wilcoxon test, function sc.tl.rank_genes_groups). Paediatric healthy and CD samples were annotated together, in order to draw direct comparisons. Marker gene expression was visualised using dot-plots where the size of the dot reflects the percentage of cells expressing the gene and color indicates relative expression.

### Epithelial cell differentiation dynamics using scVelo

Fetal epithelial cell dynamics in small bowel samples were analysed using scVelo 1.24 package implementation in Scanpy 1.4.5 (Bergen *et al.*, unpublished; Svensson and Pachter, 2018). The data was sub-clustered to epithelial cells and split into two groups 6-8 PCW (including F6.7, F6.9, F.7.9), and 9-10 PCW (including F9.9, F10, F10.1, F10.2). The clustering and visualisation was repeated using the same parameters as above for the sub-clustered cells. The data was then processed using default parameters following preprocessing as described in Scanpy scVelo implementation.

The gene-specific velocities are obtained by fitting a ratio between unspliced and spliced mRNA abundances and then computing how the observed abundances change from those observed in steady state. The ratio of ‘spliced’, ‘unspliced’, and ‘ambiguous’ transcripts was calculated to be 0.67, 0.25, 0.07, and 0.76, 0.17, 0.06 for 6-8 PCW and 9-10 PCW groups, respectively. The samples were pre-processed using functions for detection of minimum number of counts, filtering and normalisation using scv.pp.filter_and_normalise and followed by scv.pp.moments function. The gene specific velocities were then calculated using scv.tl.velocity with mode set to stochastic, and scv.tl.velocity_graph functions) and visualised using scv.pl.velocity_graph function. In addition, we used tl.recover_latent_time function to infer a shared latent time from splicing dynamics and plotted the genes along time axis sorted by expression along dynamics using scv.pl.heatmap function.

### Using receptor-ligand pairs to infer cell-cell interactions

To infer cell-cell interactions we applied the CellPhoneDB v2.0 python package (Efremova *et al.*, unpublished; Vento-Tormo *et al.*, 2018) to three separate datasets: 1) fetal cells from duo-jejunum and ileum; 2) healthy paediatric samples; and 3) Crohn’s disease samples. Log transformed and normalised counts, and cell type annotations were used as input. To narrow down to the most relevant interactions, we looked for specific interactions classified by ligand/receptor expression in more than 10% of cells within a cluster.

### Logistic regression for cellular composition and temporal classification of organoids

The python package Sklearn implementation linear_model.LogisticRegression (Pedregosa et al., 2011) was used to predict the cellular composition and temporal identity of the two organoids datasets. Expression matrix and annotation labels of all primary cells from small bowel were used as an input for training the model. We used C=0.20, solver=‘saga’ and penalty set to L1 to favour sparsity in the scRNA-seq expression matrix. The classifier estimated sparsity was 96.2% and lr.score was 0.9899. We took into account predictions with probability higher than 50% and used top labels to assign clusters. The relative abundance of predicted epithelial/mesenchymal cell-types in organoids was shown as percentage of cells per experimental condition (p1 vs p17, or RSPO vs WNT3A).

### Percentage of cells and statistical analysis

First, we calculated relative abundance of each epithelial cell type as percentage of cells per condition (control and CD, fetal ileum and fetal duodenum). We test for statistical significance using two-tailed t.test (R version 3.5.0) and report the p-values as extracted by fdrtool package (statistics = p_values). To assess regional contribution to epithelial clusters, we used total number of cells and two-way ANOVA for multiple comparisons (GraphPad).

### Common transcription factors between CD and fetal epithelium

For comparisons between fetal and paediatric epithelium, we merged and analysed cells collected from the matching anatomical location (fetal terminal ileum only) and enriched using the same strategy. First, we selected all transcription factors (TF) based on a list obtained from (http://bioinfo.life.hust.edu.cn/AnimalTFDB/#!/download) and used Scanpy sc.tl.rank_genes_groups function (Wilcoxon test) to select differentially expressed TF (from total of 1529 expressed TF) between inflamed CD (Extended Fig. 7D, arrows) versus control and non-inflamed epithelium (Extended Fig. 7D, no arrows). Out of these, we selected TFs that were differentially expressed in CD patients and upregulated in fetal epithelium and plotted their relative mean expression as a heatmap using sns.clustermap, z-score calculated for genes (rows).

### Data availability

Processed single-cell RNA sequencing objects will be available for online visualisation and download at **gutcellatlas.org.**

## Supplementary titles and legends

**Extended Figure 1: Quality control of the three datasets.** A) and B) Bar plots with a number of droplets after QC and Scrublet doublet exclusion grouped by cell type group. Scatter plots show the number of genes over the number of counts per cell, where each dot is a cell. Sample contribution to each cluster in C) Control and D) Crohn’s disease datasets. E) Bar plots with number of cells in fetal dataset and UMAP with sample contribution to each cluster. F) Number of fetal cells captured in each timepoint colored by enrichment strategy (top) and region (bottom). For fetal samples, the average cell recovery was 1,800 with a total of 62,849 cells at a mean depth of 13, 570 reads per cell and 3,027 mean genes per cell. For paediatric samples, the average recovery was 1,400 cells with 8,093 reads per cell and 1,859 mean genes per cell.

**Extended Figure 2: Cell type groups and their marker genes identified in fetal and paediatric datasets.** UMAP plots of fetal and paediatric datasets (A and C, respectively) broadly grouped into seven groups: epithelial (blue), mesenchymal (dark pink), neural (orange), endothelial (green), immune (light pink), and erythroid lineage (brown). Dot Plots of relative expression and percentage of cells expressing marker genes in fetal (B) and paediatric (D) datasets. The color bars match the cell type group colors.

**Extended Figure 3: Regional differences in epithelium of embryonic and fetal gut.** A) Number of cells of broad epithelial cell clusters (Epithelial progenitor, SI Epi, LI Epi, Secretory cells) in each of three embryonic/fetal regions (duo-jejunum, ileum and colon). B) Sub-clustered embryonic and fetal epithelial cell abundance in fetal duo-jejunum (DU) versus ileum (IL). SI Epi = small intestinal epithelium, LI Epi= large intestinal epithelium

**Extended Figure 4: Transcriptional profile of mesothelial serosa cells in embryonic and fetal human gut.** A) Embryonic/fetal gut feature plots with relative expression of epithelial, mesenchymal marker genes as well as key genes specific to serosa cells B) Signalling molecule expression in serosa cells versus FOXL1+ fibroblasts in embryonic/fetal gut.

**Extended Figure 5: Marker gene expression in fetal intestinal organoids.** A) and B) Feature plot with mesenchymal and epithelial marker gene expression in WNT3A- and WNT3A+ organoids. C) Feature plot of epithelial marker genes in p1 and p17 organoid cells.

**Extended Figure 6: Crohn’s disease patient dataset.** A) UMAP projection of Crohn’s disease samples colored by cell type annotation. B) Dot plot with relative expression and percentage of cells expressing marker genes in paediatric Crohn’s disease dataset. C) Bar plots with percentage of cells in Crohn’s disease (CD) dataset as compared with healthy paediatric (Control) dataset, grouped by the broad cell group (epithelial, stromal (mesenchymal, endothelial and glial), T cells and B cells).

**Extended Figure 7:** Dot plots with relative expression and percentage of cells expressing marker genes in paediatric A) healthy and B) Crohn’s disease epithelium. Barplots show abundance of epithelial cell subsets in C) healthy controls (n=8) and D) children with Crohn’s disease (n=7). Arrows point to samples that were grouped as “non-inflamed Crohn’s disease”.

**Supplementary Table 1:** A) Fetal single-cell sample metadata (proximal small bowel n=21,597, distal small bowel n=21,147, large bowel n=20,110) B) Paediatric single-cell sample metadata. C) Organoid single-cell sample metadata.

**Supplementary Table 2. Media composition of cultured human fetal organoids**

## References

Alimperti, S. and Andreadis, S. T. (2015) ‘CDH2 and CDH11 act as regulators of stem cell fate decisions’, Stem cell research, 14(3), pp. 270–282.

Barker, N. et al. (1999) ‘Restricted high level expression of Tcf-4 protein in intestinal and mammary gland epithelium’, The American journal of pathology, 154(1), pp. 29–35.

Barker, N. et al. (2007) ‘Identification of stem cells in small intestine and colon by marker gene Lgr5’, Nature, 449(7165), pp. 1003–1007.

Bayraktar, O. A. et al. (Nature Neuroscience, in press) ‘Single-cell in situ transcriptomic map of astrocyte cortical layer diversity’. doi: 10.1101/432104.

Becht, E. et al. (2018) ‘Dimensionality reduction for visualizing single-cell data using UMAP’, Nature biotechnology. doi: 10.1038/nbt.4314.

Bergen, V. et al. (unpublished) ‘Generalizing RNA velocity to transient cell states through dynamical modeling’. doi: 10.1101/820936.

Bettess, M. D. et al. (2005) ‘c-Myc is required for the formation of intestinal crypts but dispensable for homeostasis of the adult intestinal epithelium’, Molecular and cellular biology, 25(17), pp. 7868–7878.

Boucherat, O. et al. (2015) ‘Epithelial inactivation of Yy1 abrogates lung branching morphogenesis’, Development, 142(17), pp. 2981–2995.

Chin, A. M. et al. (2016) ‘A Dynamic WNT/β-CATENIN Signaling Environment Leads to WNT-Independent and WNT-Dependent Proliferation of Embryonic Intestinal Progenitor Cells’, Stem cell reports, 7(5), pp. 826–839.

Chin, A. M. et al. (2017) ‘Morphogenesis and maturation of the embryonic and postnatal intestine’, Seminars in cell & developmental biology, 66, pp. 81–93.

Cilieborg, M. S., Boye, M. and Sangild, P. T. (2012) ‘Bacterial colonization and gut development in preterm neonates’, Early Human Development, pp. S41–S49. doi: 10.1016/j.earlhumdev.2011.12.027.

Dupaul-Chicoine, J., Dagenais, M. and Saleh, M. (2013) ‘Crosstalk between the intestinal microbiota and the innate immune system in intestinal homeostasis and inflammatory bowel disease’, Inflammatory bowel diseases, 19(10), pp. 2227–2237.

Efremova, M. et al. (unpublished) ‘CellPhoneDB v2.0: Inferring cell-cell communication from combined expression of multi-subunit receptor-ligand complexes’. doi: 10.1101/680926.

Ellinghaus, D. et al. (2013) ‘Association between variants of PRDM1 and NDP52 and Crohn’s disease, based on exome sequencing and functional studies’, Gastroenterology, 145(2), pp. 339–347.

Fordham, R. P. et al. (2013) ‘Transplantation of expanded fetal intestinal progenitors contributes to colon regeneration after injury’, Cell stem cell, 13(6), pp. 734–744.

Fukamachi, H. et al. (2001) ‘Mesenchymal transcription factor Fkh6 is essential for the development and differentiation of parietal cells’, Biochemical and biophysical research communications, 280(4), pp. 1069–1076.

Gao, S. et al. (2018) ‘Publisher Correction: Tracing the temporal-spatial transcriptome landscapes of the human fetal digestive tract using single-cell RNA-sequencing’, Nature cell biology, 20(10), p. 1227.

Geske, M. J. et al. (2008) ‘Fgf9 signaling regulates small intestinal elongation and mesenchymal development’, Development, 135(17), pp. 2959–2968.

Grosse, A. S. et al. (2011) ‘Cell dynamics in fetal intestinal epithelium: implications for intestinal growth and morphogenesis’, Development, 138(20), pp. 4423–4432.

Guiu, J. et al. (2019) ‘Tracing the origin of adult intestinal stem cells’, Nature, 570(7759), pp. 107–111.

Harper, J. et al. (2011) ‘The transcriptional repressor Blimp1/Prdm1 regulates postnatal reprogramming of intestinal enterocytes’, Proceedings of the National Academy of Sciences of the United States of America, 108(26), pp. 10585–10590.

Ito, K. et al. (2014) ‘Gene targeting study reveals unexpected expression of brain-expressed X-linked 2 in endocrine and tissue stem/progenitor cells in mice’, The Journal of biological chemistry, 289(43), pp. 29892–29911.

Kaestner, K. H. et al. (1996) ‘Clustered arrangement of winged helix genes fkh-6 and MFH-1: possible implications for mesoderm development’, Development, 122(6), pp. 1751–1758.

Kaestner, K. H. et al. (1997) ‘The mesenchymal winged helix transcription factor Fkh6 is required for the control of gastrointestinal proliferation and differentiation’, Genes & development, 11(12), pp. 1583–1595.

Kaestner, K. H. (2019) ‘The Intestinal Stem Cell Niche: A Central Role for Foxl1-Expressing Subepithelial Telocytes’, Cellular and molecular gastroenterology and hepatology, 8(1), pp. 111–117.

Karlsson, L. et al. (2000) ‘Abnormal gastrointestinal development in PDGF-A and PDGFR-(alpha) deficient mice implicates a novel mesenchymal structure with putative instructive properties in villus morphogenesis’, Development, 127(16), pp. 3457–3466.

Kinchen, J. et al. (2018) ‘Structural Remodeling of the Human Colonic Mesenchyme in Inflammatory Bowel Disease’, Cell, 175(2), pp. 372–386.e17.

Korinek, V. et al. (1998) ‘Depletion of epithelial stem-cell compartments in the small intestine of mice lacking Tcf-4’, Nature genetics, 19(4), pp. 379–383.

Kraiczy, J. et al. (2016) ‘Assessing DNA methylation in the developing human intestinal epithelium: potential link to inflammatory bowel disease’, Mucosal immunology, 9(3), pp. 647–658.

Kraiczy, J. et al. (2019) ‘DNA methylation defines regional identity of human intestinal epithelial organoids and undergoes dynamic changes during development’, Gut, 68(1), pp. 49–61.

Kurahashi, M. et al. (2008) ‘Platelet-derived growth factor signals play critical roles in differentiation of longitudinal smooth muscle cells in mouse embryonic gut’, Neurogastroenterology and motility: the official journal of the European Gastrointestinal Motility Society, 20(5), pp. 521–531.

Li, N. et al. (2019) ‘Memory CD4 T cells are generated in the human fetal intestine’, Nature immunology, 20(3), pp. 301–312.

Lin, C.-C. et al. (2014) ‘Bhlhe40 controls cytokine production by T cells and is essential for pathogenicity in autoimmune neuroinflammation’, Nature Communications. doi: 10.1038/ncomms4551.

Madison, B. B. et al. (2005) ‘Epithelial hedgehog signals pattern the intestinal crypt-villus axis’, Development, 132(2), pp. 279–289.

Mallow, E. B. et al. (1996) ‘Human enteric defensins. Gene structure and developmental expression’, The Journal of biological chemistry, 271(8), pp. 4038–4045.

Matthews, S. M. et al. (2019) ‘The Crohn’s disease associated SNP rs6651252 impacts MYC gene expression in human colonic epithelial cells’, PLOS ONE, p. e0212850. doi: 10.1371/journal.pone.0212850.

Muncan, V. et al. (2011) ‘Blimp1 regulates the transition of neonatal to adult intestinal epithelium’, Nature communications, 2, p. 452.

Mustata, R. C. et al. (2013) ‘Identification of Lgr5-independent spheroid-generating progenitors of the mouse fetal intestinal epithelium’, Cell reports, 5(2), pp. 421–432.

Nowotschin, S. et al. (2019) ‘The emergent landscape of the mouse gut endoderm at single-cell resolution’, Nature, 569(7756), pp. 361–367.

Parikh, K. et al. (2019) ‘Colonic epithelial cell diversity in health and inflammatory bowel disease’, Nature, 567(7746), pp. 49–55.

Polański, K. et al. (2019) ‘BBKNN: Fast Batch Alignment of Single Cell Transcriptomes’, Bioinformatics. doi: 10.1093/bioinformatics/btz625.

Sato, T., Stange, D. E., et al. (2011) ‘Long-term Expansion of Epithelial Organoids From Human Colon, Adenoma, Adenocarcinoma, and Barrett’s Epithelium’, Gastroenterology, pp. 1762–1772. doi: 10.1053/j.gastro.2011.07.050.

Sato, T., van Es, J. H., et al. (2011) ‘Paneth cells constitute the niche for Lgr5 stem cells in intestinal crypts’, Nature, 469(7330), pp. 415–418.

Schreurs, R. R. C. E. et al. (2019) ‘Human Fetal TNF-α-Cytokine-Producing CD4 Effector Memory T Cells Promote Intestinal Development and Mediate Inflammation Early in Life’, Immunity, 50(2), pp. 462–476.e8.

Scialdone, A. et al. (2015) ‘Computational assignment of cell-cycle stage from single-cell transcriptome data’, Methods, 85, pp. 54–61.

Scikit-learn: Machine Learning in Python, Pedregosa et al., JMLR 12, pp. 2825–2830, 2011.

Shoshkes-Carmel, M. et al. (2018) ‘Subepithelial telocytes are an important source of Wnts that supports intestinal crypts’, Nature, 557(7704), pp. 242–246.

Shyer, A. E. et al. (2015) ‘Bending gradients: how the intestinal stem cell gets its home’, Cell, 161(3), pp. 569–580.

Smillie, C. S. et al. (2019) ‘Intra- and Inter-cellular Rewiring of the Human Colon during Ulcerative Colitis’, Cell, 178(3), pp. 714–730.e22.

Sonntag, B. et al. (2007) ‘Preterm birth but not mode of delivery is associated with an increased risk of developing inflammatory bowel disease later in life’, Inflammatory Bowel Diseases, pp. 1385–1390. doi: 10.1002/ibd.20206.

Svensson, V. and Pachter, L. (2018) ‘RNA Velocity: Molecular Kinetics from Single-Cell RNA-Seq’, Molecular Cell, pp. 7–9. doi: 10.1016/j.molcel.2018.09.026.

Teng, Y.-S. et al. (2020) ‘Upexpression of BHLHE40 in gastric epithelial cells increases CXCL12 production through interaction with p-STAT3 in Helicobacter pylori-associated gastritis’, FASEB journal: official publication of the Federation of American Societies for Experimental Biology, 34(1), pp. 1169–1181.

Traag, V. A., Waltman, L. and van Eck, N. J. (2019) ‘From Louvain to Leiden: guaranteeing well-connected communities’, Scientific Reports. doi: 10.1038/s41598-019-41695-z.

Vento-Tormo, R. et al. (2018) ‘Single-cell reconstruction of the early maternal-fetal interface in humans’, Nature, 563(7731), pp. 347–353.

Walton, K. D. et al. (2012) ‘Hedgehog-responsive mesenchymal clusters direct patterning and emergence of intestinal villi’, Proceedings of the National Academy of Sciences of the United States of America, 109(39), pp. 15817–15822.

Walton, K. D., Freddo, A. M., et al. (2016) ‘Generation of intestinal surface: an absorbing tale’, Development, pp. 2261–2272. doi: 10.1242/dev.135400.

Walton, K. D., Whidden, M., et al. (2016) ‘Villification in the mouse: Bmp signals control intestinal villus patterning’, Development, 143(3), pp. 427–436.

Wehkamp, J. et al. (2007) ‘The Paneth Cell α-Defensin Deficiency of Ileal Crohn’s Disease Is Linked to Wnt/Tcf-4’, The Journal of Immunology, pp. 3109–3118. doi: 10.4049/jimmunol.179.5.3109.

Wolf, F. A., Angerer, P. and Theis, F. J. (2018) ‘SCANPY: large-scale single-cell gene expression data analysis’, Genome biology, 19(1), p. 15.

Wolock, S. L., Lopez, R. and Klein, A. M. (2019) ‘Scrublet: Computational Identification of Cell Doublets in Single-Cell Transcriptomic Data’, Cell systems, 8(4), pp. 281–291.e9.

Yanai, H. et al. (2017) ‘Intestinal stem cells contribute to the maturation of the neonatal small intestine and colon independently of digestive activity’, Scientific reports, 7(1), p. 9891.

Ye, H. et al. (1997) ‘Hepatocyte nuclear factor 3/fork head homolog 11 is expressed in proliferating epithelial and mesenchymal cells of embryonic and adult tissues’, Molecular and cellular biology, 17(3), pp. 1626–1641.

Yu, F. et al. (2018) ‘The transcription factor Bhlhe40 is a switch of inflammatory versus antiinflammatory Th1 cell fate determination’, The Journal of experimental medicine, 215(7), pp. 1813–1821.

Yui, S. et al. (2018) ‘YAP/TAZ-Dependent Reprogramming of Colonic Epithelium Links ECM Remodeling to Tissue Regeneration’, Cell stem cell, 22(1), pp. 35–49.e7.

Zhang, C. et al. (2019) ‘YY1 mediates TGF-β1-induced EMT and pro-fibrogenesis in alveolar epithelial cells’, Respiratory research, 20(1), p. 249.

