## Supplementary figures and images for "Single-cell sequencing of developing human gut reveals transcriptional links to childhood Crohn’s disease"

### Supplemental Figures

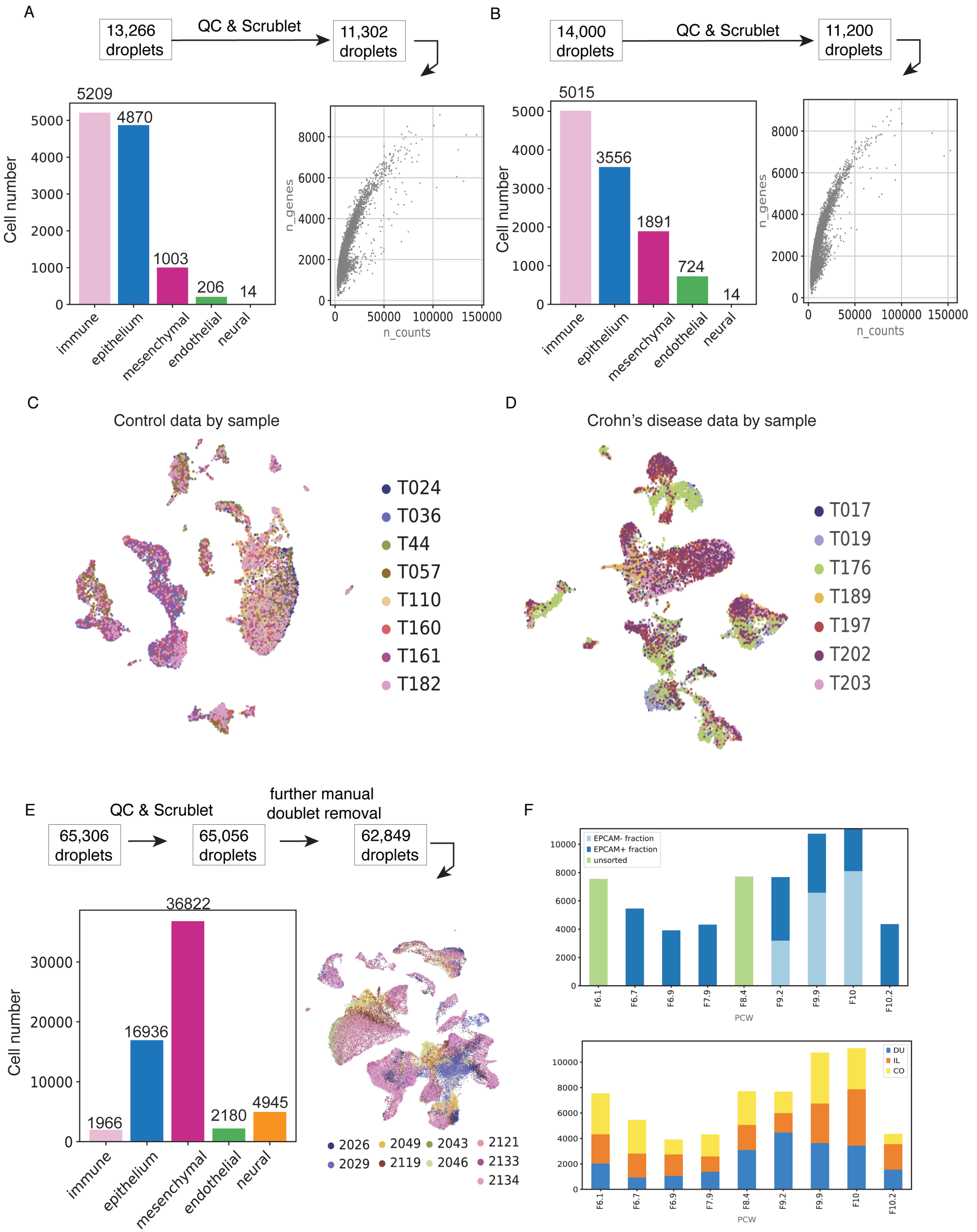

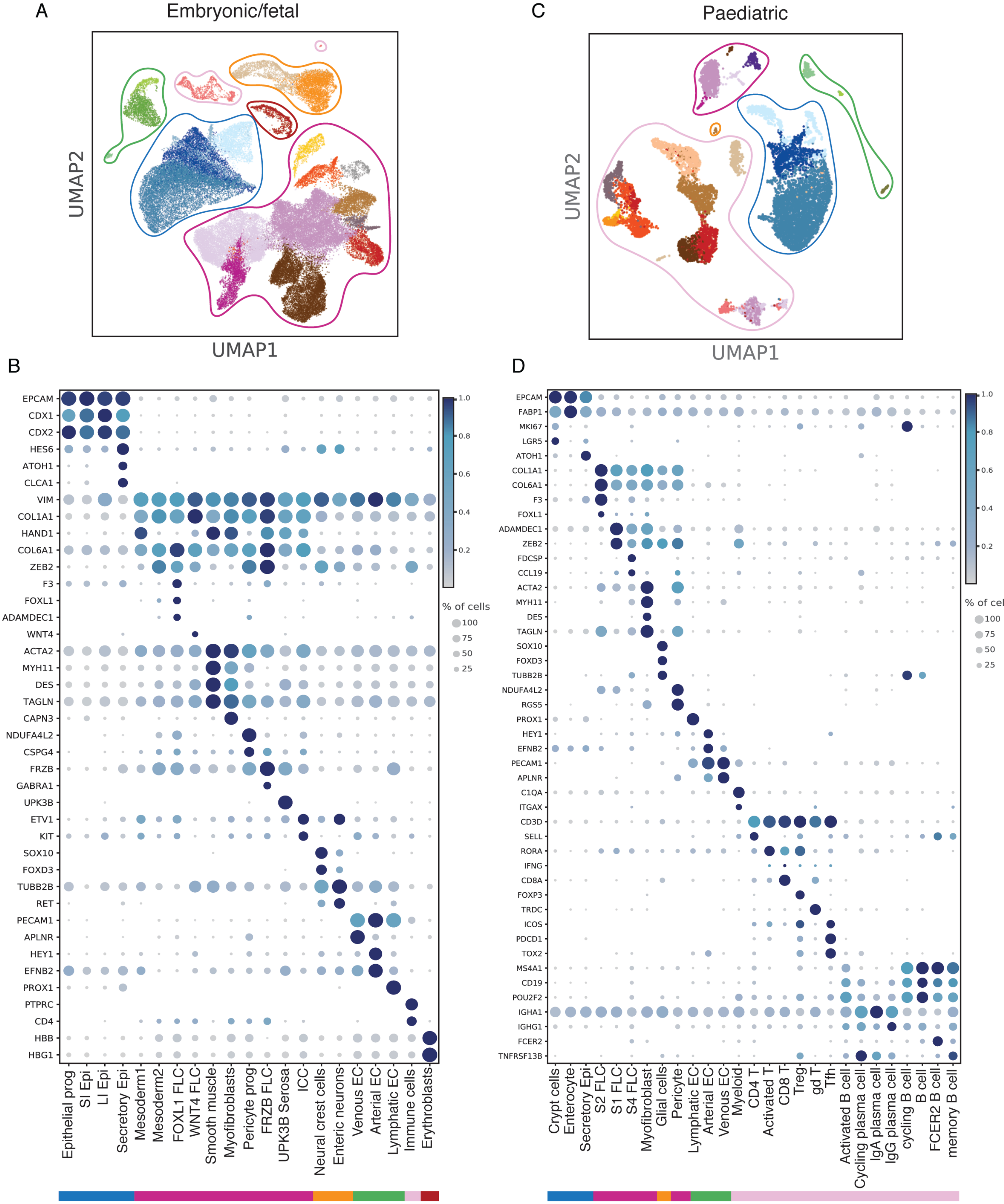

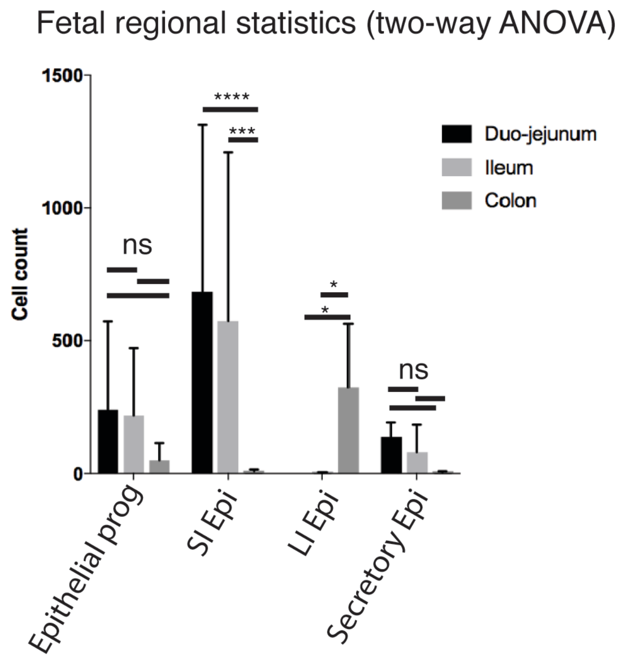

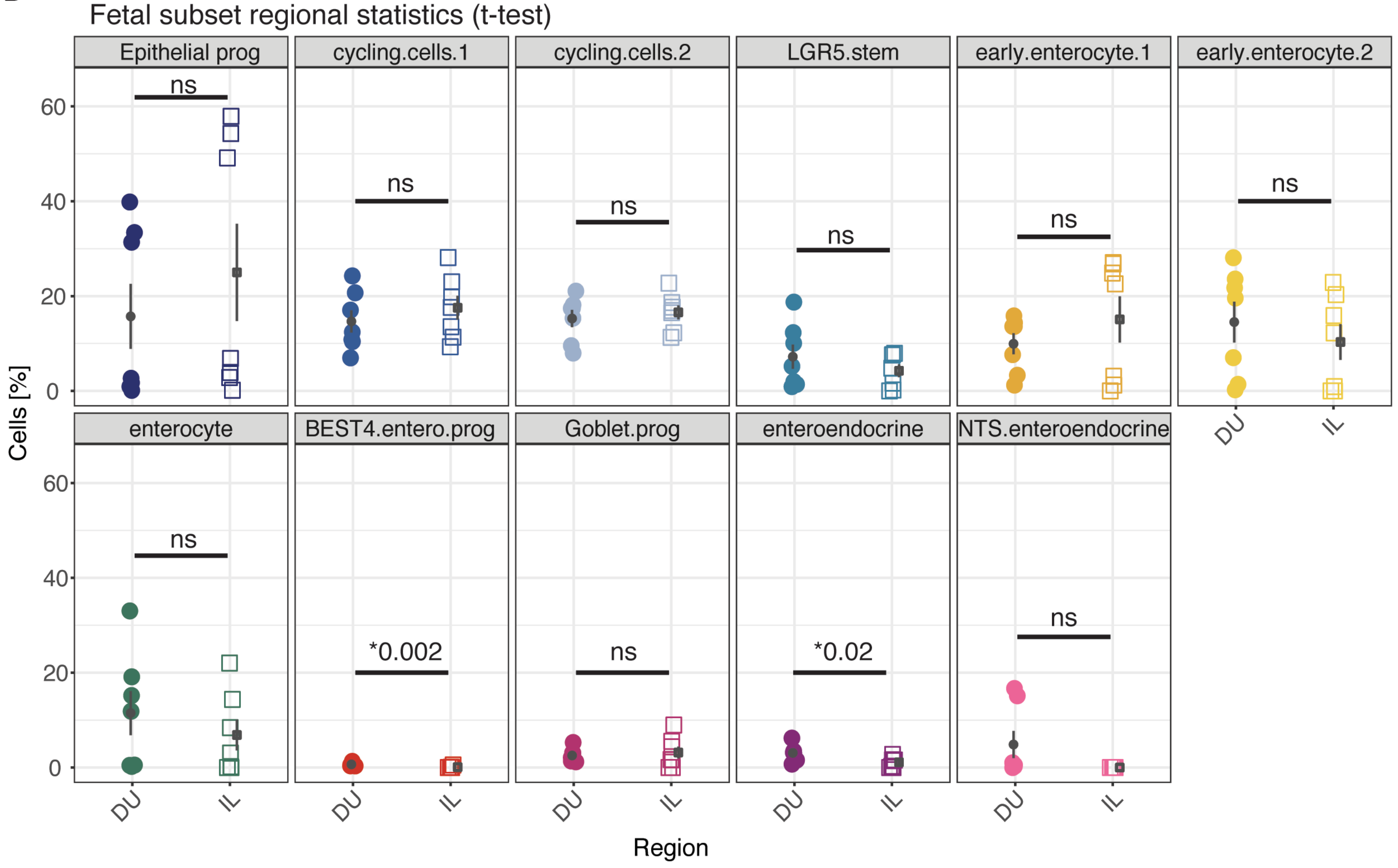

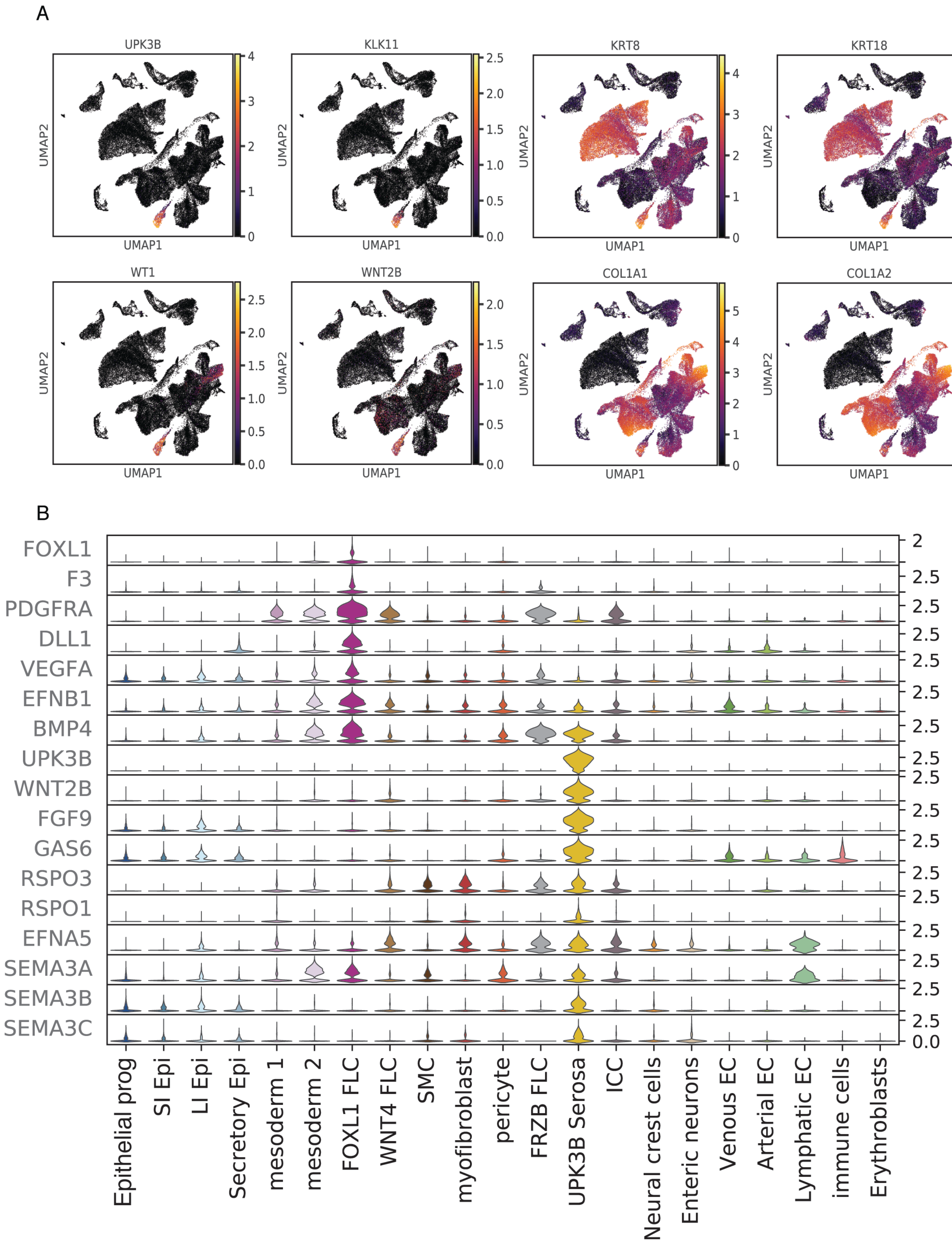

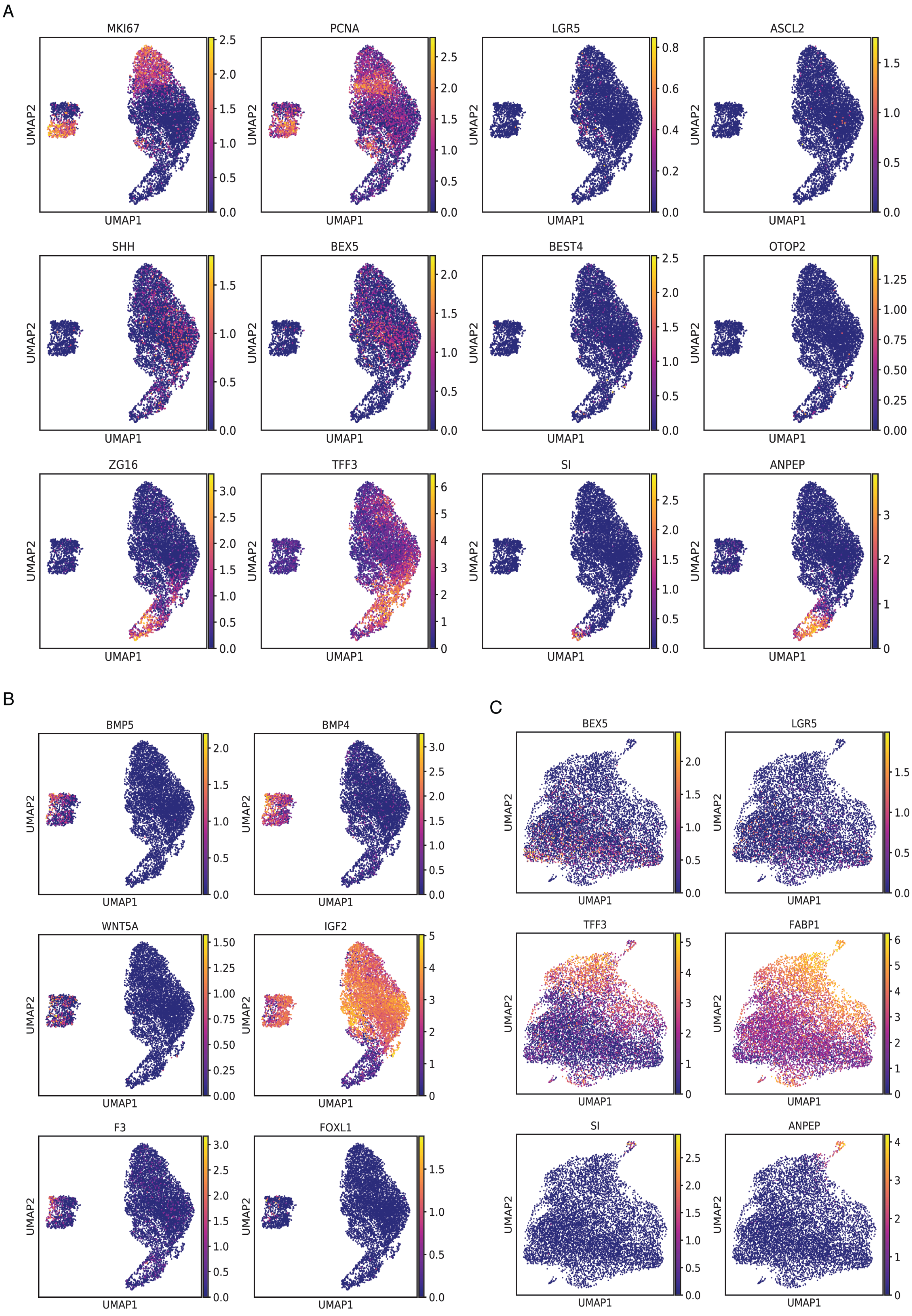

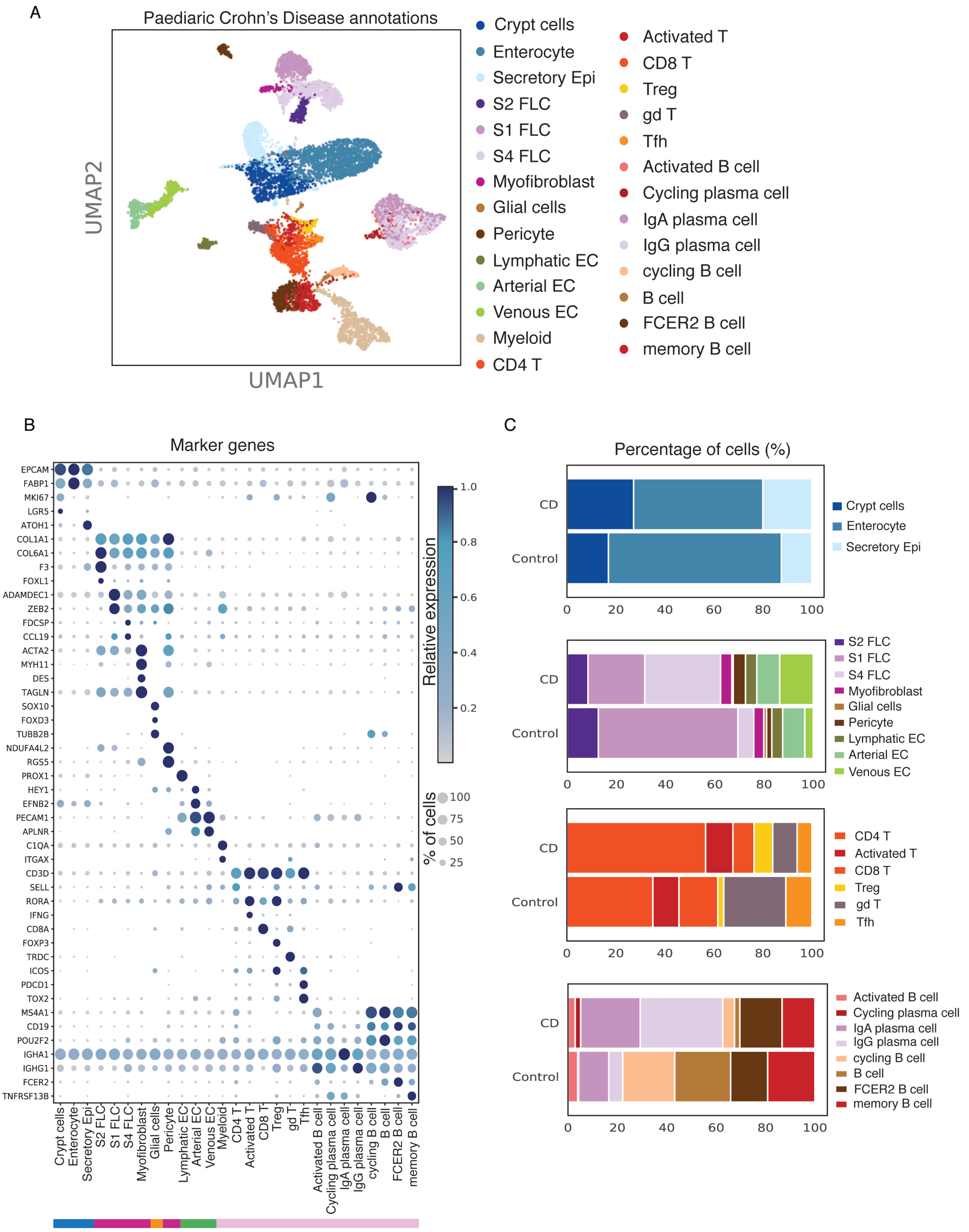

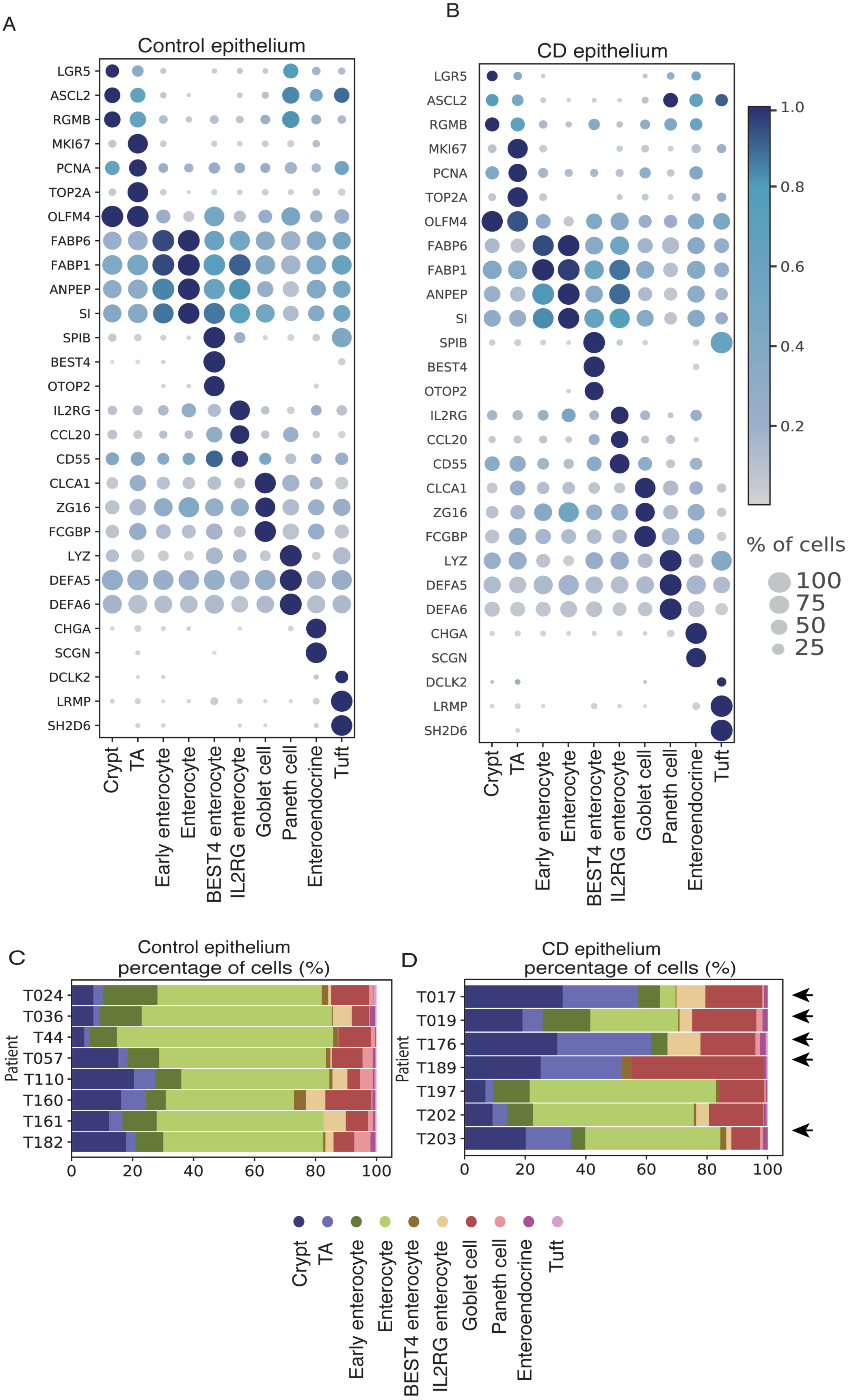
